# Amplification and delay of auditory feedback for walking: effects on variability, cadence, and temporal cortex activity

**DOI:** 10.64898/2026.01.24.700725

**Authors:** Jingxian Gu, Presley Honeyman, Yi Gao, Dobromir Dotov

## Abstract

When we walk, our footsteps generate rich acoustic information about foot timing, dragging, loading rate, etc. The role of self-generated or endogenous auditory feedback for gait control remains underexplored. We investigated how delay and amplification of self-produced footstep sounds modulate gait. Thirty healthy young adults walked overground while receiving real-time lateralized playback of the sound produced by their feet. Amplified auditory feedback (AAF) was delivered with no, shorter, or longer delay, and either at full or half amplification. Cortical hemodynamic responses were recorded concurrently in prefrontal and temporal regions of interest (ROI) with mobile functional near-infrared spectroscopy (fNIRS). Spatiotemporal gait parameters were analyzed as change relative to baseline. We found that amplification without delay reduced variability by almost 10%, consistent with strengthened sensorimotor coupling via enhanced perceptual access to foot-ground interaction dynamics. Delay increased cadence instead of reducing it. At the level of hemodynamic response, the ROI containing the superior temporal gyrus (STG) exhibited an increase during walking relative to quiet standing, especially in the condition with no delay and full amplification. To explain the effects of amplification and delay on gait timing, we offer two alternative models, one using anticipatory dynamic systems and the other Bayesian predictive processing.

## Delay and amplification of auditory feedback for walking: effects on variability, cadence, and temporal cortex activity

When we walk, the sounds of our footsteps reach our ears, but we rarely notice them. Our actions in the environment often produce sounds that provide a rich source of information. Sounds also provide information about other agents in the world. One implication of this fact is that animals must be able to distinguish between self-generated and other-generated sounds. This is a critical aspect of survival skills for animals in the wild who rely on the sense of hearing to monitor for danger. This has motivated the hypothesis that downstream connections from motor to auditory areas in the brain allow perceptual processes to anticipate self-generated sounds and thus separate them from other-generated sounds (Schneider & Mooney, 2018). Searching for this proposed mechanism for selectivity has led to the discovery of a cortical filter in mice (Schneider et al., 2018). In humans, evidence suggests that neural oscillations in response to auditory stimuli differentiate sounds from self-produced actions (Ross et al., 2017). Connections between auditory and motor cortices are strong in humans (Merchant et al., 2015). While cumulative evidence supports a connection from motor to auditory perceptual processes, it is less clear whether auditory information is automatically involved in gait control.

The openness of the auditory system and its propensity to couple with motor control have been exploited to create sensory augmentation strategies to improve movement. Experimental approaches to gait rehabilitation have been exploiting *external* or *exogenous* cueing to improve gait. Clinically, sensory augmentation or sensory cues have been widely used in gait rehabilitation, with evidence supporting their benefit for improving gait performance (Ploughman et al., 2018), and auditory cueing has been one of the commonly employed strategies (Baker et al., 2007; Thaut et al., 1996). Numerous studies suggest that auditory cues have a positive influence on gait patterns (Ready et al., 2022; Shin & Chung, 2022). Specifically, rhythmic auditory cueing has been shown to enhance gait by providing continuous external timing signals that facilitate movement coordination in different clinical populations such as those with Parkinson’s disease or stroke survivors (Dvorsky et al., 2011; Ploughman et al., 2018). Repeated tones or music with a set auditory rhythm support functional task performance in individuals with Parkinson’s disease (Rochester et al., 2005; Spaulding et al., 2013).

Another promising avenue for improving and retraining gait is to emphasize feedback for self-produced action, understood as *internal*, *endogenous*, or *self-cueing*. Feedback for gait retraining can take multiple forms (Spencer et al., 2021). Among them, continuous movement sonification during treadmill walking enhances proprioception (Hermann et al., 2011; Schmitz et al., 2014; Wall et al., 2024) and improves performance (Pang, Cheng, et al., 2025). Manipulating the spectral properties of sonified footstep sounds affects gait and perceived body weight (D’Adamo et al., 2026; Tajadura-Jiménez et al., 2015). Despite its relevance to gait improvement and retraining, amplified auditory feedback (AAF) from self-produced footsteps has been investigated far less than exogenous cueing. Here, we examined how non-instructed AAF affected gait. This was motivated by the idea that humans, like other animals, monitor self-produced footstep sounds and should therefore automatically respond to AAF.

Interestingly, sensory feedback can be beneficial not only when it is amplified but also when it is delayed, as revealed by the effects of delayed-auditory feedback (DAF) on speech. Even though DAF increases dysfluencies in people without speech difficulties (Stuart et al., 2002) and reduces speech rate (Hashimoto & Sakai, 2003), paradoxically, it can improve speech in people who stutter (Kalinowski & Stuart, 1996; Stuart et al., 2004). In the latter case, it is possible that DAF serves as an additional timing cue that bypasses faulty internal timing circuits. Aside from speech, DAF has been studied experimentally in finger tapping paradigms via its effects on mean negative asynchrony (MNA), the amount of time by which tapping tends to precede the target event. The consequences of delayed feedback on tapping include an additive combination of tactile-kinesthetic and auditory streams (Aschersleben & Prinz, 1997). Also in tapping, there is a motor state-dependency where the response is a function of finger trajectory at the time of DAF onset (Pfordresher & Dalla Bella, 2011). DAF for walking has rarely been investigated. Menzer et al. (2010) explored a large range of delays, from zero to more than a full gait cycle, and found a sinusoidal pattern of increase and reduction of the stride interval as a function of the delay. They also reported a similar pattern of reduction in the sense of agency, meaning that for some delays the sensory stimulation was experienced as externally caused.

We manipulated the delay and amplitude in real-time using a custom-built apparatus for delayed auditory feedback for walking (AAF). Across different trials, the feedback was delayed by different amounts or not, and the sound level was amplified or not, or masked with broadband noise. Our main hypotheses were that: 1) Augmenting self-produced footstep sound would have a positive effect on gait, specifically by reducing variability and increasing speed; 2) Self-produced footstep sound would entrain gait, specifically by reducing cadence in the delayed auditory feedback conditions.

We also used mobile brain imaging to conduct an exploratory investigation of cortical activation patterns associated with the different feedback conditions. We used functional near-infrared spectroscopy (fNIRS), a mobile brain imaging modality that is suitable for experimental paradigms with free movement that aim to bridge the gap between neuroscience of sensory-motor integration and whole-body tasks (Yücel et al., 2021). We focused on three regions of interest (ROI) that were theoretically motivated and were accessible by fNIRS, which has low penetration. The dorsolateral prefrontal cortex (dlPFC) was considered because of its role in divided attention. The dlPFC exhibits increased activation in dual tasks such as walking with a concurrent cognitive task (Rahman et al., 2021), walking on a cluttered terrain with a cognitive task (Krelove et al., 2025), and walking while performing an auditory Stroop task (Kvist et al., 2023, 2026). An increase in dlPFC in this study would be consistent if the feedback acted as a distracting novel task. We also focused on the premotor and supplementary motor areas (PM/SMA) because of their role in sequencing and planning motor actions. Increased activation has been observed with fNIRS while walking with exogenous rhythmic auditory cueing (Vitorio et al., 2018). The third area of interest was the superior temporal gyrus (STG). It is part of auditory-motor integration pathways, primarily studied in speech but also a candidate for gait-related corollary discharge pathways targeting sensory processing (Reznik et al., 2014; Schneider et al., 2018; van Elk et al., 2014). Importantly, the STG is not as deep as the primary and associative auditory areas, which are difficult to reach with fNIRS.

## Methods

### Participants

Thirty healthy young adults (*N*=30, Male/Female = 18/12, age range 20-33) were recruited for this study. Participants were screened prior to testing and included based on the following criteria: ability to walk independently for one hour, no diagnosis of a neurological condition, free from musculoskeletal problems, not taking medication that could impact motor performance, and no hearing problems. Participants received monetary compensation. Ethics approval was obtained from the University of Nebraska Medical Center IRB (#0794-22-EP).

### Procedure

Participants signed informed consent forms prior to the start of testing. Participants were asked to provide their height, weight, and demographic information, including gender and age, and to complete a questionnaire about self-reporting any hearing or musculoskeletal issues. The researcher introduced the experimental procedure to the participants and adjusted the delayed auditory feedback apparatus to each participant’s preferred amplitude. The researcher then fitted the participant with the experimental apparatus, including the functional near-infrared spectroscopy (fNIRS) system, the inertial measurement units (IMUs), and the delayed feedback device. After all apparatus were in place, the researcher checked the fNIRS signal and calibrated both the brain imaging and movement kinematics systems. The researcher introduced the walking route and allowed the participants to practice walking for one loop. Baseline gait parameters were taken during the first 30 seconds of a pre-test trial, during which participants walked at their self-selected pace. Prior to data collection, participants’ head dimensions were measured from the nasion (Nz) to the inion (Iz) and between the left and right preauricular points (LPA-RPA) to select the best-fitting cap. Before each recording, the researcher inspected signal quality and calibrated the fNIRS system.

### Apparatus

The apparatus for delayed auditory feedback during walking (AAF) was custom-built. It captured a two-channel sound stream using microphones worn on the shoes and played it back in real-time through headphones. The system included two microphones, one mounted on each shoe, a mobile computing device worn on the belt, and headphones (Sony WH-1000XM4, Sony, Tokyo, Japan), see Figure 1A-B.

**Figure 1.**
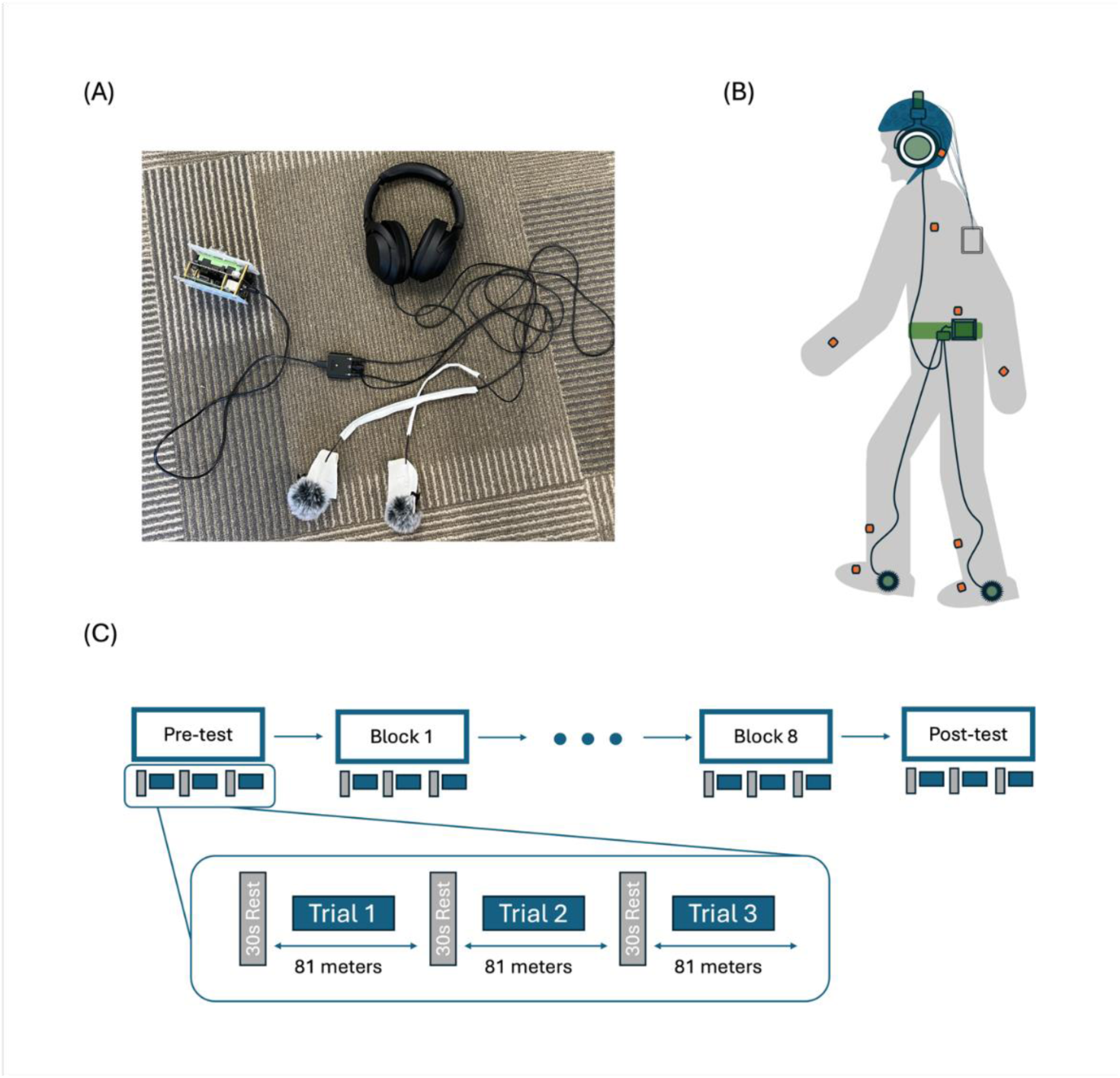
Delayed Auditory Feedback for Walking (AAF) apparatus and procedures. (A) Custom-built apparatus consisting of two microphones, a Raspberry Pi, and headphones. (B) Illustration of a participant wearing the apparatus. Orange blocks represent the APDM sensors; the blue cap represents the fNIRS system with optical sensors; the green headphones with microphones and cables represent the custom-built apparatus used to deliver footstep sounds. (C) Experimental design consisting of a Pre-Test, eight blocks, and a Post-Test. Each block includes three trials.

The microphones were omnidirectional lavalier microphones with windproof hairballs. They were mounted on the dorsal lateral side of the shoe facing down. To prevent noise from friction between the microphone and the shoe, we used a thermoplastic plate to create a concave tray-like structure that isolated the microphone from the shoe while securely fixing it in place with medical tape. To minimize latency, the microphones were connected to the computer via cables and a two-channel interface (RODE AI-Micro, RØDE, Sydney, Australia). We measured a 15-millisecond latency of the apparatus from microphone to headphones. The computer was a Raspberry Pi 5 (Raspberry Pi Foundation, UK) running a GNU/Linux operating system and a custom Python program. The program introduced a combination of delay, amplification, and/or noise masking depending on the experimental condition.

Individual levels of natural and amplified footstep sounds were not measured due to the complexity of reliably measuring in-headphone sound levels while the participant was walking. We estimated the range of amplification using a sealed sound level meter and the range of self-selected sound levels. The amplification from the microphone at the foot level to the headphones was between −3.48 and .32 dBA at full amplification (−9.48 and −5.68 dBA at half amplification). To put this in the context of natural sounds, we calculated the following relationship between natural ambient and amplified sound levels, considering how sound level decreases with distance from the foot to the head. For a person whose ears were 160 cm from the ground, making footsteps that were 68.6 dBA at the level of the microphone placed 10 cm from the ground, the level of the peak ambient footstep sound would be 44.52 dBA, while the amplified sound inside the headphone at median sound level setting would be 66.3 dBA (60.3 dBA at half amplification). For reference, peak sound levels of adult humans walking indoors on various apartment floors have been measured in the range from 31.5 to 45.5 dBA (Park & Lee, 2017), although the literature on walking sound intensity is surprisingly sparse. With respect to the constant masking noise condition, the sound level at the median setting would be 55.7 dBA (49.7 at half amplitude). Background noise from fans and computers in the testing area was 45 dBA. The testing area had a flat synthetic walking surface optimized for gait experiments. Participants were asked to arrive for the data collection session with their comfortable athletic shoes.

For gait kinematic measures such as cadence, participants wore inertial measurement units (Opal IMU’s, APDM Wearable Technologies, Portland, OR, USA), placed using elastic straps on the left and right feet, left and right shanks, lumbar area, chest, left and right wrists, and forehead. For mobile brain imaging, participants wore an fNIRS system (NIRSport 2, NIRx, Berlin, Germany) consisting of a data logging device on a back strap and a cap with sensors. Participants wore an fNIRS cap with optodes while the portable battery-powered data-acquisition hardware was secured using a back-mounted strap. The fNIRS apparatus included an accelerometer used to record head movement and assist with correction of motion-related artifacts.

### Design

There were a total of 10 walking conditions, including Pre-test and Post-test. In all conditions, participants were instructed to walk at a comfortable pace overground. The walking surface was a VCT (vinyl composition tile) floor, and they wore their own comfortable shoes. In the Pre-test and Post-test, participants wore the headphones, but no sound was played. We manipulated delay at three levels (no, low, high) and amplitude at two levels (full or half amplification). Low delay was 12.5% of the participant’s preferred step duration. High delay was 25% of the participant’s preferred step duration, or a phase-offset of 90 degrees. Amplitude was self-determined based on the participant’s comfort level before experimental trials. The masking condition consisted of pink (broadband) noise played either at full or half amplification level. The combination of these factors resulted in the following conditions: (a) Delay No, Amplitude Low, (b) Delay No, Amplitude High, (c) Delay Low, Amplitude Low, (d) Delay Low, Amplitude High, (e) Delay High, Amplitude Low, (f) Delay High, Amplitude High, (g) Masking Noise Low, (h) Masking Noise High.

A block of three trials was completed in each condition. Each trial consisted of two loops around the track, amounting to 81 meters total. Between trials, there was a rest period of 30 seconds, or more if participants requested. Each trial included 15 seconds of quiet standing before walking. This design was necessary to provide the trials’ baseline for fNIRS analysis. Trials were blocked by condition, which was also necessary for fNIRS analysis. The order of conditions was randomized across participants, save for Pre-test and Post-test, which were first and last, respectively. Between blocks (conditions), participants rested freely until they were ready to begin the next block, see Figure 1C.

### Measures

#### Kinematics

We used the proprietary algorithms of the sensor manufacturer’s software suite, Mobility Lab (APDM, Portland, OR, USA), to obtain spatiotemporal gait parameters. Only straight walking portions of the trial were used for these measures; the algorithm automatically detects turning and excludes steps. Cadence and speed were measured in each gait cycle, step to step. Stride length was measured in each stride. We combined the left and right sides into a single trial measure and retained the trial-average of left-side and right-side step/stride parameters. For variability of cadence, speed, and stride length, the coefficient of variation (*CV*) was calculated separately for the left and right sides and then averaged across left and right.

#### Cortical hemodynamic response

The probes consisted of 16 sources and 15 detectors operating at wavelengths of 760 nm and 850 nm, sampled at 6.7 Hz. This yielded 34 long measurement channels and 8 short-separation channels. We used the fNIRS optodes’ location decider (fOLD) (Zimeo Morais et al., 2018) to design the probe montage on an EEG 10-5 cap and provide coverage of cortical regions relevant to the experimental task, including dlPFC, PM/SMA, and STG. The three ROI-level responses were weighted averages of channel-level responses. The spatial arrangement of sources, detectors, and channels is shown in Appendix Table A1, and ROI’s and corresponding channels in Appendix Figure A1. Areas with low sensitivity, such as the primary and secondary auditory cortex, were excluded from analysis.

The preprocessing of fNIRS data was done in Homer3 (Huppert et al., 2009). Continuous recordings were segmented according to the experimental conditions. Each segment included 15s of data preceding condition onset and 30 s following condition offset to retain sufficient baseline and post-condition information. First, channels with inadequate signal quality were identified and excluded using the *hmrR_PruneChannels* function with a signal-to-noise ratio criterion of 6.7. Raw light-intensity measurements were subsequently converted to changes in optical density using *hmrR_Intensity2OD*. Motion artifacts were identified on a channel-by-channel basis using *hmrMotionArtifactByChannel* function with window length 30 s, and duration 5 s. Motion contaminated signals were subsequently corrected using spline interpolation (*hmrR_MotionCorrectSpline*) with the spline parameter p=0.99, followed by wavelet-based motion correction (hmrR_MotionCorrectWavelet) with an interquartile-range parameter of 1.5. Optical-density signals were then band-pass filtered between 0.01 and 0.2 Hz. Accelerometer signals were filtered between 0.01 and 0.5 Hz. Following preprocessing, optical-density changes were converted to concentration changes in oxygenated hemoglobin (ΔHbO) and deoxygenated hemoglobin (ΔHbR) using the modified Beer-Lambert law. We focused the analysis on HbO because it has been reported to have preferable properties (Kinder et al., 2022; Wilson et al., 2014). The differential pathlength factor was set to 1.0 for both wavelengths.

Condition related changes in HbO concentrations were estimated using a general linear model (GLM) implemented in Homer3. The hemodynamic response function (HRF) was modeled using a Gaussian basis function over a time window of 0 to 45 s relative to condition onset. To account for superficial hemodynamic signals, short-separation measurements with source-detector distances ≤ 15 mm were included as nuisance regressors. For long-separation measurements, the short-separation channel exhibiting the greatest temporal correlation with that measurement was selected for short-separation regression (flagSSmethod=1). Slow temporal drift was modeled using a third-order polynomial.

### Statistical Analysis

We used linear mixed-effects models for statistical analysis using the *lme4* package in R (Bates, 2010). This is a flexible approach that allows unbalanced design matrices and is more tolerant relative to some of the assumptions of traditional methods (Raudenbush & Bryk, 2002).

## Results

### Kinematics

To facilitate interpretation, we expressed all measures as change (Δ%) relative to the average of the three pre-test trials. Because conditions were repeated observations over consecutive blocks, the independent variables were coded in the design matrix of the linear models as time-varying dummy variables: delay∈ {0,1,2}, masking noise∈ {0,1}, amplitude∈ {0,1,2}. For the amplitude dummy variable, 0 corresponded to no feedback amplification in the Post-test trials. We included block order as a predictor to account for potential time effects such as fatigue. An incremental approach to the statistical model was followed (Singer & Willett, 2003). We started with an overall intercept model and included all random effects that the model could support numerically and without overfitting. There was an a priori reason to expect a nonlinear effect of delay when going from zero to small (eight of the step duration) to large (a quarter of the step) delay because of the periodic nature of footstep delay. Therefore we included a squared delay term in the linear model. The largest possible model was specified as follows: Y*_ij_*=β_0_+σ_0*i*_+σ_1*i*_+β_1_BlockOrder*_ij_*+β_2_Masking*_ij_*+σ_3*i*_+β_3_Delay*_ij_*+β_4_Delay^2^*_ij_*+σ_5*i*_+β_5_Amplitude*_ij_*+β_6_Amplitude*_ij_*Delay*_ij_*+σ*_ij_* where *i* is participant, *j* is trial, σ_0*i*_ is individual baseline variability, σ_0*i*_ is individual baseline variability, σ_1*i*_ is individual dependent on block number, σ_3*i*_ is individual dependence on delay, and σ*_ij_* is the residual variability. The summary statistics of dependent variables^1^ per condition are in Appendix Table B1.

### Δ Cadence

Block number had a positive effect (β = 0.29442, *SE* = 0.05599, *p* < .001, η²_partial_ = 0.47) with cadence increasing over successive blocks of trials throughout the duration of the experiment. Delay showed a U-shaped relationship with Δ cadence: the linear term was negative (β = −1.18696, *SE* = 0.22074, *p* < .001, η²_partial_ = 0.05), but the quadratic term was positive (β = 0.86939, *SE* = 0.10022, p < .001, η²_partial_ = 0.09). This means that the shorter delay tended to reduce cadence, but the higher delay increased it, see Figure 2A. Neither masking noise (β = 0.08792, *SE* = 0.11369, *p* = 0.4396, η²_partial_ =8.03e-04) nor amplitude (β = −0.08766, *SE* = 0.07835, *p* = 0.2714, η² partial = 0.04) had statistically significant effects, although amplitude showed a slight negative trend (See Figure 2A).

**Figure 2.**
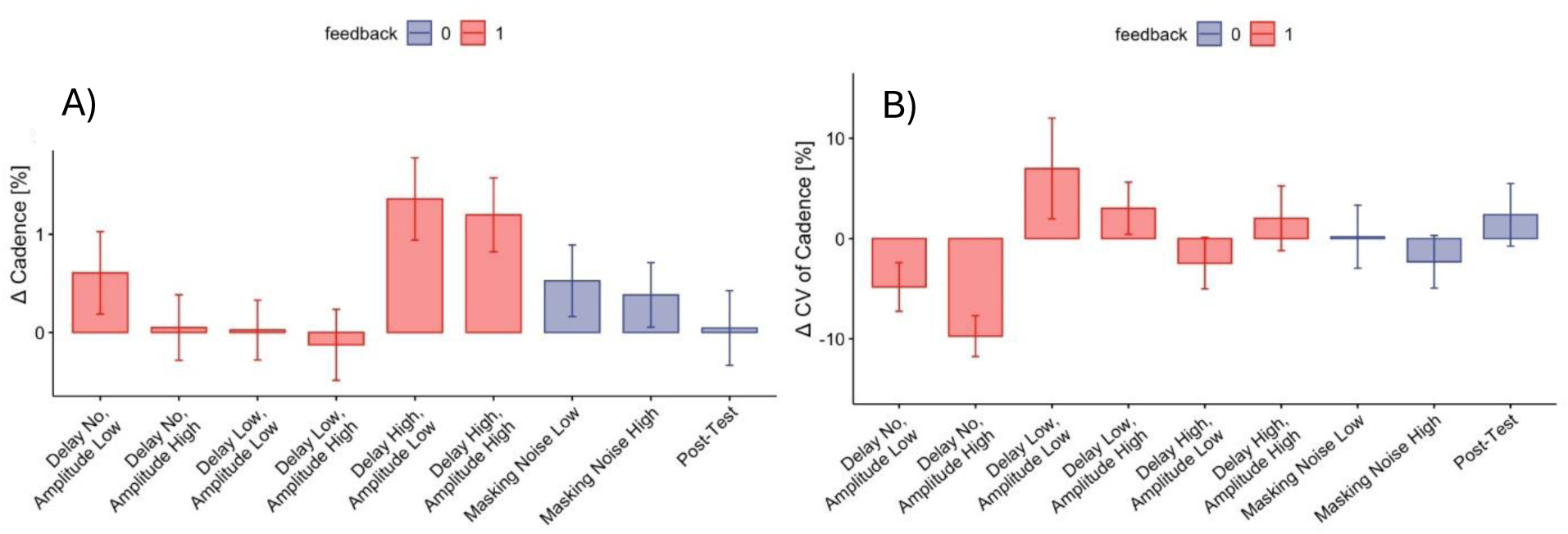
Change in spatiotemporal gait parameters in each walking condition (*M*±*SE*) relative to the baseline. The serial trend along blocks was subtracted by using the block order term in the fitted statistical model. A) Cadence. B) Variability.

### Δ Speed

The significant model for change in speed included block number, masking noise, and delay, see Appendix Table C2 and Appendix Figure D1. The effect of block was statistically significant, with participants showing an increase in gait speed over successive blocks of trials (β = 0.8276, *SE* = 0.1358, *p* < .001, η²_partial_ =0.56). Delay showed a significant positive effect (β = 0.6419, *SE* = 0.1723, *p* < 0.01, η²_partial_ = 0.27), suggesting that increasing delay was associated with higher gait speed. Masking noise had no significant effect on change of speed (β = 0.3352, *SE* = 0.2258, *p* = 0.138065, η²_partial_ = 2.92e-03).

### Δ Stride length

The selected model included block number, masking noise, delay and amplitude, see Appendix Table C3 and Appendix Figure D1. The effect of block was statistically significant, with participants showing a decrease in stride length over time (β = 0.48690, *SE* = 0.08536, *p* < .001, η²_partial_ = 0.52). Delay had a U-shaped relationship with Δ stride length: the linear term was positive (β = 1.19559, *SE* = 0.28635, *p* < .001, η²_partial_ = 0.03), but the quadratic term was negative (β = −0.56639, *SE* = 0.13223, p < .001, η²_partial_ = 0.02). The effect of masking noise (β =0.14313, *SE* = 0.14990, *p* = 0.33996) and amplitude (β = 0.10533, *SE* = 0.10696, *p* = 0.33097) were not statistically significant.

### Δ CV of cadence

The final model included effects for block number, masking noise, delay, amplitude, a quadratic effect of delay, and an interaction between delay and amplitude, see Appendix Table C4. Together, the effects of delay and amplitude suggested a strong negative effect in the ‘Delay No, Amplitude High’ condition, see Figure 2B. The statistical model showed that block had a decreasing trend, but the effect was not statistically significant (*p = 0.057*). Masking noise had a positive effect, suggesting an increase in variability (β = 5.7941, *SE* = 2.8297, *p* < 0.05, η²_partial_ = 5.43e-03). The linear term of delay had a positive effect (β = 14.3156, *SE* = 5.8819, *p* < 0.05, η²_partial_ = 7.62e-03), and the quadratic term had a negative effect (β = −8.8971, *SE* = 2.4496, *p* < .001, η²_partial_ = 0.02).

Amplitude had a significant negative effect (β = −5.8851, *SE* = 1.8901, *p* <0.05 η²_partial_ = 0.01). Also, the interaction between delay and amplitude was significant (β =4.6118, *SE* = 2.1636, *p* < .05, η²_partial_ =5.90e-03). To test in which conditions variability changed relative to baseline, we conducted an additional test where all conditions were treated as levels of a categorical factor and Pre-Test was the reference level. We found that the ‘Delay No, Amplitude High’ condition was significantly lower compared to the Pre-test (β = −9.4574, *SE* = 4.6317, *p* < 0.05, *Cohen’s d* = −0.3458801), all other conditions (*p* > .05).

### Cortical activation

We estimated changes in cortical response compared to the pre-trial quiet standing baseline. The pre-test was informative here, given the relative novelty of mobile brain imaging in walking. We used a statistical model with a single categorical factor in which each walking condition was a level, and pre-test served as the reference condition: Y*_ij_* = β_0_ + σ_0*i*_ + β_1_BlockOrder*_ij_* + σ_1*i*_ + β_2_WalkingCondition*_ij_* + σ*_ij_*, where *i* is participant, *j* is block, σ_0*i*_ is variability of the individual average intercept, σ_1*i*_ is variability of the individual dependence on block order, and σ*_ij_* is the residual variability. The same model was fitted separately for each ROI without false discovery rate correction. The summary statistics are in Appendix Table E.

Within the *dorsolateral Prefrontal Cortex (dlPFC)*, neither the pre-test nor any of the other walking conditions were significantly different from zero (all *p*’s>.05), indicating that, on average the dlPFC did not exhibit a change in activation during walking relative to standing, see Figure 3A. Expectedly, block order was not a significant factor (*p*>.05). Within the *Premotor and Supplementary Motor Area (PM/SMA)*, the pre-test was not significantly different from zero (*p*>.05). Despite a negative trend, the ‘Delay High, Amplitude Low’ condition was significantly lower than the pre-test, (β = −0.000071, *SE* = 0.000031, *p* < .05, ω²_partial_ = 0.02), whereas other walking conditions were not different than zero either (all *p*’s>.05, Figure 3B). Block order was not a significant factor (*p*>.05). Within the *Superior Temporal Gyrus (STG)*, the pre-test was significantly higher from zero (β = −0.000073, *SE* = 0.000029, *p* < .05, ω²_partial_ = 0.06). Most other conditions were not different from the pre-test (all *p*’s>.05), implying that they were also higher than zero. The ‘Delay No, Amplitude High’ condition was significantly higher than the pre-test (β = 0.000094, *SE* = 0.000033, *p* < .01, ω²_partial_ = 0.03), see Figure 3C. Block order was not a significant factor (*p*>.05).

**Figure 3.**
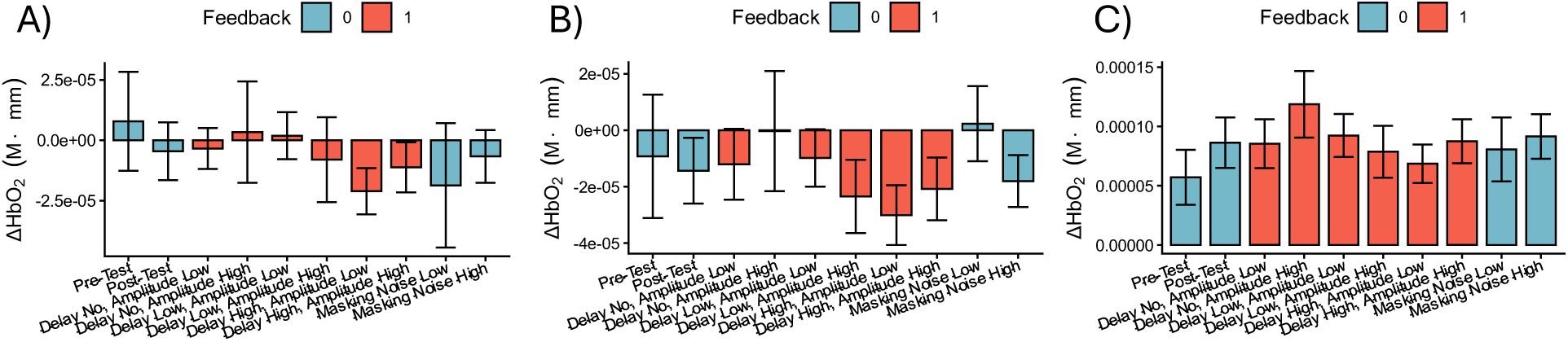
Mean hemodynamic response ΔHbO_2_ per condition (*M*±*SE*) in each of the three ROI’s: A) dorsolateral Prefrontal, B) Premotor and Supplementary Motor, C) Superior Temporal Gyrus.

## Discussion

In this study, altered AAF for walking resulted in several effects on gait parameters without participants being instructed to synchronize with or correct the feedback. This confirms our overall hypothesis that the human brain is poised to exploit all available information, including the acoustic cues, to control motor performance in the environment. Our three main findings concerned reduction of variability, increase of cadence, and relatively little importance of masking. The reduction of variability with non-delayed but amplified self-generated footstep sounds is an expected and beneficial consequence of emphasizing salient interactions between the walker and the environment. This is consistent with observations that combining discrete and continuous information can lead to stronger effects of feedback (Pang, Feltham, et al., 2025). Foot interaction with the ground is a complex mechanical process, far from being a discrete point-like event. It comprises striking, sliding, rotating, dragging, and unloading lower leg segments, all of which dissipate energy in the form of a highly structured acoustic signal.

Using closed headphones does not eliminate, and may even emphasize, bone conduction as a form of covert auditory feedback with zero delay. For this reason, the results should be interpreted in terms of multisensory integration rather than pure presence or absence of feedback. Indeed, sensory redundancy should be seen as the norm because other sensory modalities such as mechano-tactile, vestibular, and visual play critical roles during walking (Roytman et al., 2025). To compare emphasized and reduced auditory feedback, we used a masking condition with a broadband noise. We found little evidence that masking changed gait, consistent with a previous study (Cornwell et al., 2020). The combined results of amplified and masked auditory feedback reveal that the brain mixes sensory modalities in a non-linear fashion: when a non-essential modality provides the opportunity for augmented information, this can improve performance, but when a non-essential modality is removed, the other ones are sufficient to maintain performance.

### Hemodynamic responses

The dlPFC did not exhibit a differential hemodynamic response in walking trials relative to the quiet standing baseline. This ROI was motivated by its role in dual task paradigms with external stimuli that serve as a secondary task in addition to walking. It appears that the auditory feedback in this study did not instantiate a distractor strong enough to elicit effects indicative of split attention. With respect to PM/SMA, there was a tendency for lower than baseline activation in most conditions, although the effect was significant only in the ‘Delay High, Amplification Low’ condition. The trend toward a decrease hemodynamic response in PM/SMA during walking relative to quiet standing may seem surprising, but it has been reported in fNIRS studies before (Koenraadt et al., 2014; Perrey, 2014). It can be explained by lower cortical demands during steady-state walking, that is, when the walker has settled into a comfortable repetitive motor pattern.

An interesting pattern we observed in this exploratory study was that the hemodynamic response in the STG was consistently higher during walking relative to quiet standing. This effect was even stronger in the ‘Delay No, Amplitude High’ feedback condition and can be considered the purest and strongest form of amplified feedback in this study. If supported in future hypothesis-driven studies, this response to amplified feedback for walking would be an exciting result which has not been reported yet, to the best of our knowledge. While STG is known to be involved in auditory-motor integration in speech (Hickok & Poeppel, 2007), less is known about its role in gait control. In a finger tapping study, STG showed increased BOLD signal during trials with self-produced versus passively listened to piano keys sounds (Reznik et al., 2014). In postural control, auditory feedback for postural fluctuations increased activation in cortical areas, including the temporal region (Pirini et al., 2011). In an fNIRS dance study, STG exhibited increased activity relative to rest (Tachibana et al., 2011).

There are theoretical reasons why STG is expected to be involved in the coupling of motor reafference and auditory feedback while walking. As stated, human and animal studies point to higher auditory areas as candidate sites for converging motor re-afference and auditory feedback pathways (Schneider & Mooney, 2018). STG also participates in a larger loop engaged in rhythmic skills which are relevant given the inherent rhythmicity of walking (J. L. Chen et al., 2009; Grahn & Rowe, 2009; Zatorre et al., 2007). Subregions of STG also respond to spectrotemporal patterns (Hickok & Poeppel, 2007; Nourski et al., 2009; Obleser et al., 2007). This has been established in speech processing but is also relevant in the present context because of the complex nature of footstep sounds, including, for example, low frequencies from ground impact and high frequencies from subsequent friction against the ground. Finally, parts of the STG and surrounding areas may respond to action sounds, or doable sounds, when integrating multiple sensory modalities (Rauschecker & Scott, 2009), which is particularly relevant when hearing one’s own footsteps. Importantly, these theoretical strands concern specific subregions of the STG and adjacent areas such as auditory areas, within the Sylvian fissure. In contrast, our analysis was defined by spatially crude and superficial ROI’s. For higher spatial resolution, future work needs to use advanced methods such as deeper brain imaging and mapping of individual brain morphology.

### Anticipatory dynamical systems and Bayesian perspectives on variability and frequency adaptation with delayed self-feedback

Feedback amplification was found to reduce inter-stride interval variability in this study, which was an expected effect. The unexpected finding was that the large delay of quarter step period was associated with an increase in cadence, contrary to our hypothesis based on previous research in speech (Yates, 1963). The only comparable study in gait also reported an increase in cadence but at a larger delay and in short walking trials (Menzer et al., 2010). In the following, we consider two mathematically formulated explanations. They are qualitative and do not have parameters with easy physiological interpretation. They also belong to alternative theoretical lineages, although our intention is not to treat them as mutually exclusive (Large et al., 2023).

#### Anticipatory synchronization, variability, and frequency adaptation from delayed self-feedback

The remarkable formal discovery of strong anticipation is that in certain dynamic systems with delayed feedback, the driven part can run ahead of the driver and thus anticipate it without storing an internal model (Dubois, 2003). In anticipatory delayed feedback master-slave systems, here referred to as leader-follower, the follower’s coupling with the leader includes self-feedback that is delayed. The idea of anticipation without prediction is counterintuitive, but it makes sense when considering that it works in tandem with synchronization. If the follower falls in a synchronized state with the leader, meaning that both leader and follower run on the same manifold, then the state of follower will be running ahead of the leader by an amount compensating for its self-feedback delay (Voss, 2000). Strong anticipation is an elegant account of delayed feedback in motor control and sensory-motor integration (Stepp, 2009; Stepp & Turvey, 2015; Voss & Stepp, 2016). It explains anticipatory phenomena in auditory-motor synchronization, including negative mean asynchrony relative to a rhythmic stimulus (Harding et al., 2025; Roman et al., 2019), interpersonal synchronization (Heggli et al., 2019; Washburn et al., 2015), and force production (Grover et al., 2021).

For a minimal model of anticipatory synchronization in oscillatory systems, the well-known Kuramoto system can be re-formulated with a self-feedback delay (Demos et al., 2019; Mirasso et al., 2017). The leader is a phase oscillator running at a fixed frequency ω*_L_*, see Eq. 1. The follower is a phase oscillator running at a frequency that is the sum of its intrinsic frequency ω*_F_* and a coupling term with the classic Kuramoto form, the difference being that the self part of the coupling is delayed by an amount τ:

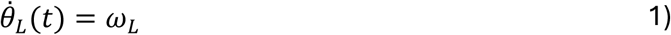

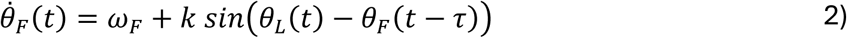

We re-formulated this model^2^ to account for the specific features of the present study. In the walking task, the environment was inert but it generated sensory stimulation such as optic flow and mechanical feedback relative to the movement of the person walking. Importantly, the walker could adapt to the room, but the room could not adapt to the walker. For this reason, in the leader-follower scheme, we treated the environment as the leader and the walker as the follower, while the frequency of the environment at any instant of time was the instantaneous frequency of the gait cycle, Eq. 3. The gait cycle was coupled to the leader via a Kuramoto-style term in which the self part was delayed by an amount τ. A second important assumption reflected the multi-modality of sensory feedback. Interaction with the environment while walking involved at least two sources of feedback: non-delayed coupling via mechanical and optical sensory modalities and coupling via amplified auditory feedback with experimentally controlled self-delay.

Therefore, the follower had two coupling terms with different self-delays, *τ_F,1_* and *τ_F,2_*, and coupling strength parameters, *k_1_* and *k_2_*, see Eq. 4. For generality, coupling with the leader could be delayed too, *τ_L_*. Finally, we added frequency adaptation, Eq. 5, (Ermentrout, 1991; Taylor et al., 2010). Frequency adaptation ran on a slower time scale μ with the tendency α to return to its intrinsic frequency Ω*_F_*, and received the same coupling as the phase but with different coupling strengths, ε*_1_* and ε*_2_*.

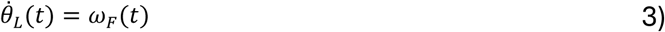

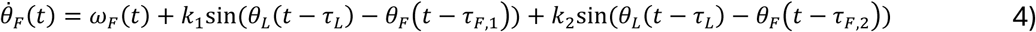

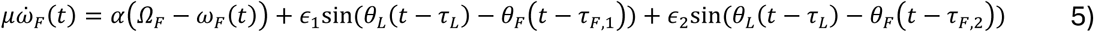

As Figure 4 shows, the model exhibited anticipatory synchronization in the form of stable negative relative phase, combined with an increase in cadence over the duration of the trial. We explored how this model accounted for variability and cadence by systematically varying the two parameters of the second self-feedback delay term: delay τ*_F_*_,2_=[0,2π] *rad* and coupling strength *k_2_*= [0,4], with ε*_2_*=2*k_2_*. We used Runge-Kutta fourth-order integration with a time step *dt*=.001 and a trial duration of 120 seconds. The time scale separation parameter was μ=10. Without loss of generality, we set α=0 because this term had a minor effect in short trials. The initial frequency was Ω*_F_*=4π rad/s, corresponding to 2 cycles per second or 120 steps per minute. The initial relative phase between leader and follower was θ*_L_*_,0_-θ*_F_*_,0_=0 *rad*. The leader coupling delay was τ*_L_*=π/6 *rad*. The parameters of the first feedback term were also fixed: coupling strength, *k_1_*=2, ε*_1_*=4, and self-delay τ*_F_*_,1_=π/5 *rad*.

**Figure 4.**
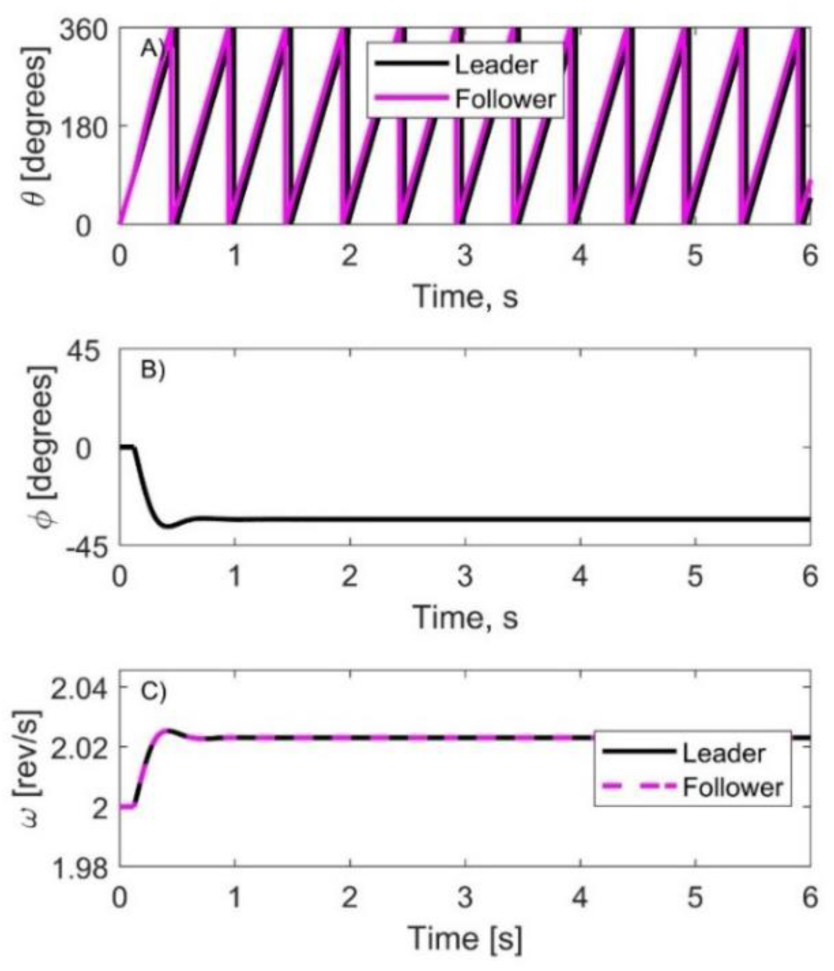
A) phase, B) relative phase, and C) frequency adaptation of the anticipatory synchronization model, Eqs. 3-5, with a small delay τ*_F_*_,2_=π/6 *rad* and an initial condition of φ=0°.

Figure 5 shows how frequency changed in the parameter space of self-delay τ*_F_*_,2_ and coupling strength *k_2_*. Corresponding to the present study, Figure 5B shows that frequency increased with increasing coupling strength when delay was fixed at τ_follower,2_=90°. Furthermore, the effect on frequency was periodic as it switched from positive to negative in the range τ_follower,2_>180°, see Figure 5C. The periodic character is consistent with the Menzer et al. (2010) study, although the exact delay values at which they observed maximum and minimum cadence may not align with the predictions of this model. Notably, the participants in their study were primed to attend to their footsteps by a recurrent self-agency questionnaire, and the trials were short. This highlights the difference between automatic and voluntary control of walking.

**Figure 5.**
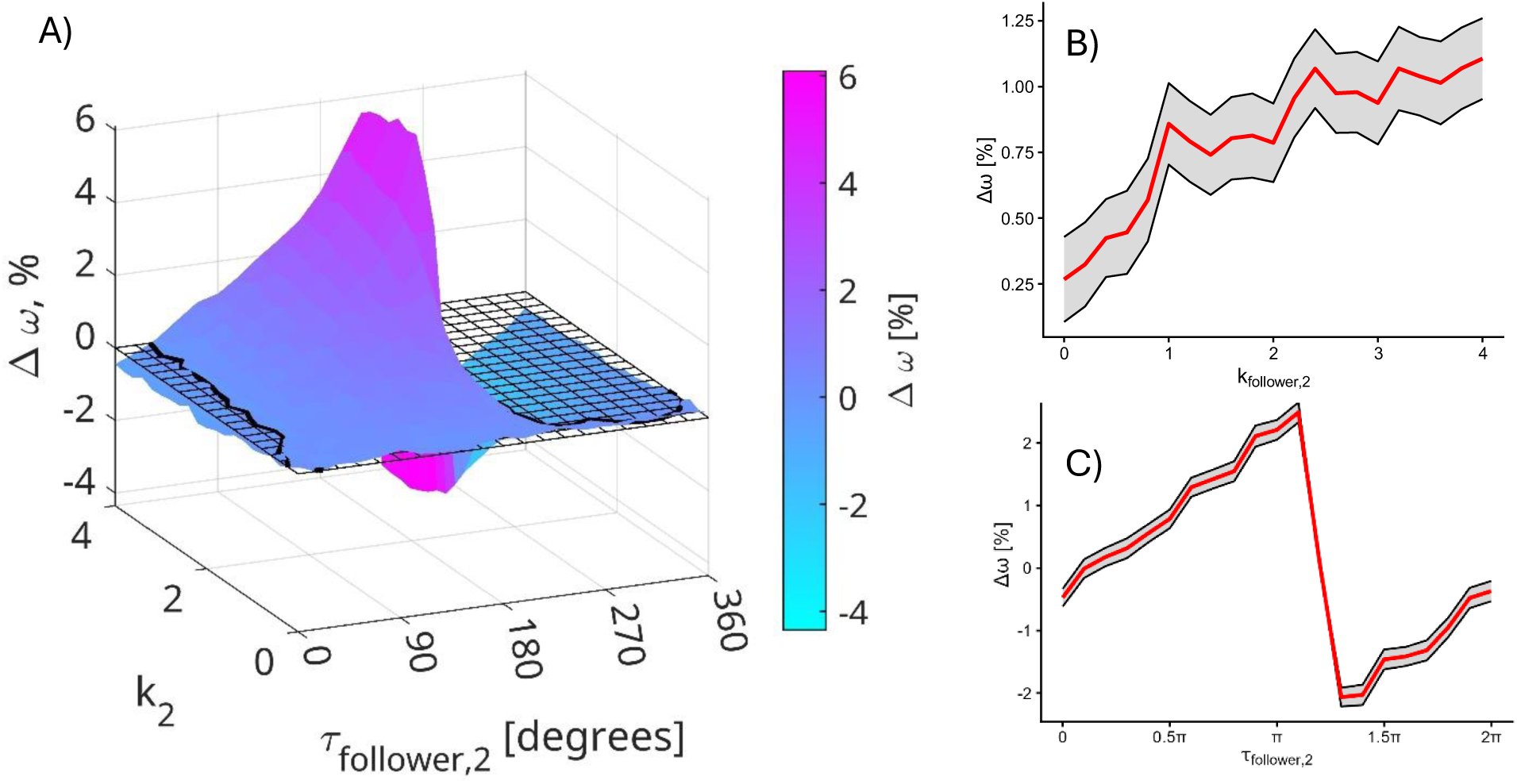
A) Frequency as a function of self-feedback strength and delay in the anticipatory dynamical model. B) Frequency adaptation when delay was fixed at τ_follower,2_=90° and strength *k*_follower,2_ was varied. *M*±*CI* over 500 simulations. C) Frequency adaptation when feedback strength was fixed at *k*_follower,2_=2 and delay τ_follower,2_ was varied. *M*±*CI* over 500 simulations.

Figure 6 shows how variability changed in the parameter space. Corresponding to the present study, when feedback was not delayed, τ_follower,2_=0°, variability decreased with increasing coupling strength, Figure 6B. Delay increased the variability of oscillation frequency, particularly at specific values, see Figure 6C, which also leads to interesting, testable predictions for future studies.

**Figure 6.**
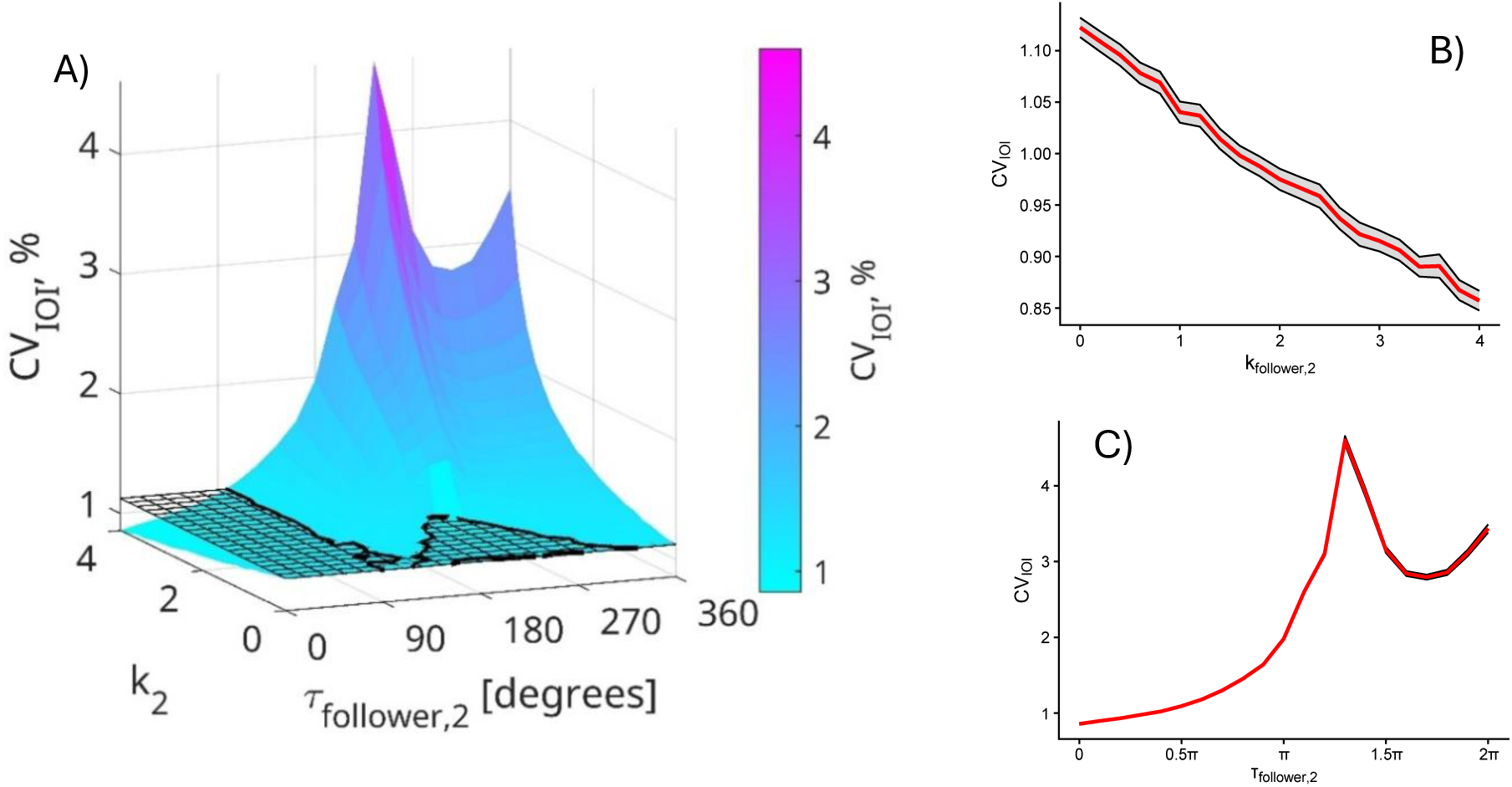
A) Oscillator interval variability as a function of self-feedback strength and delay in the anticipatory dynamical model. The grid denotes baseline. B) *CV*_IOI_ when delay was fixed at τ_follower,2_=0° and strength k_follower,2_ was varied. *M*±*CI* over 500 simulations. C) *CV*_IOI_ when feedback strength was fixed at *k*_follower,2_=4 and delay τ_follower,2_ was varied. *M*±*CI* over 500 simulations.

#### Bayesian predictive coding and multisensory causal inference

A previous study interpreted delayed footstep feedback in terms of the relationship between sensorimotor mismatch and sensorimotor prediction (Menzer et al., 2010). Effects of sensory delay in motor control have traditionally been explained by internal models (Körding & Wolpert, 2004; Wolpert et al., 1995). In this approach, a forward model uses motor commands and current state estimates to predict interactions with the environment, while an error-correction process updates this state estimate based on the difference between expected and actual sensory feedback. This theory addresses a key result in sensory-motor tapping synchronization: negative mean asynchrony (NMA). NMA is the tendency for motor events to precede target sensory stimuli. In tapping studies, reducing sensory information leads to a larger NMA by increasing temporal uncertainty of the target (Aschersleben, 2002; Steen & Keller, 2013). Arguably, because the delay inherent in sensory processing makes it impossible to verify that action matches the timing of the target stimulus, the predictive mechanism compensates by driving the action early. This style of theoretical explanation can be expanded in the form of predictive coding to account for synchronization with complex rhythms, not just interval tapping (Vuust & Witek, 2014). Predictive processing for rhythmic sensory-motor synchronization has been given a concrete Bayesian expression (Cannon, 2021). Here, we propose a proof-of-principle model that demonstrates how the two key findings can arise from a compact Bayesian perception-action architecture: reliability-weighted causal integration stabilizes cadence under non-delayed amplified feedback, whereas delayed feedback produces nonlinear cadence adaptation when causal attribution decreases and late actions are penalized.

Delayed auditory feedback during walking can be interpreted as temporally distorted sensory evidence about the current gait cycle. Critically, because footstep sounds are embedded in a continuous multimodal perception-action loop, delayed auditory feedback may not always be treated as evidence from the current step. We therefore implemented a minimal Bayesian causal-inference toy model to illustrate how delayed and amplified auditory feedback could affect cadence and cadence variability^3^. The model estimates the current step period, *T*, by combining an internal prediction, undelayed natural gait-related sensory evidence, and experimentally amplified auditory evidence. Amplification increases the precision of auditory evidence, consistent with reliability-weighted cue integration (Ernst & Banks, 2002). This is achieved by *σ_aud_*(*A*) = *σ*_0_ + *σ_g_*/(1 + *γA*), where *A* is the experimentally controlled auditory amplification, *σ*_0_ is an irreducible auditory timing-noise floor, *σ_g_* is the amplitude-sensitive component of auditory noise, and *γ* scales how strongly amplification improves precision. The corresponding observation variance is 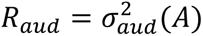. In contrast, delay reduces the inferred probability that the auditory feedback and current step share a common cause, consistent with Bayesian causal inference in multisensory perception (Körding et al., 2007; Rohe & Noppeney, 2015). The causal attribution term was defined as

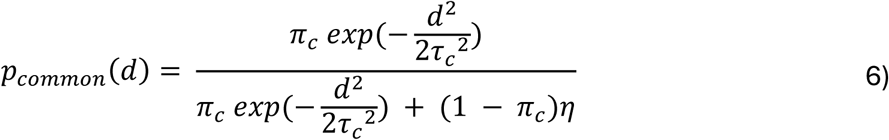

Here, *p_common_*(*d*) is the posterior probability that the amplified auditory feedback and the current step have a common cause; *d* is the auditory delay expressed as a fraction of the current step period; *π_c_* is the prior probability of a common cause before considering the observed delay; *τ_c_* is the temporal binding width, with smaller values making common-cause attribution decrease more rapidly with delay; and *η* is the relative likelihood of the separate-cause model. When delay is small, the common-cause probability is high and auditory feedback is likely to be attributed to the current step. When delay is larger, the common-cause probability decreases and the auditory cue is increasingly treated as a separate event.

The final sensory estimate was computed by Bayesian model averaging between a fused natural-plus-auditory estimate and a natural-evidence-only estimate:

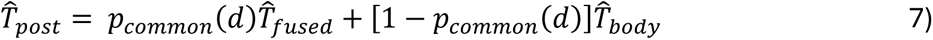

Here, *T̂_post_* is the final posterior estimate of the current step period; *T̂_fused_* is the precision-weighted estimate obtained by combining natural gait-related sensory evidence with amplified auditory evidence. *T̂_body_* is the estimate based on natural gait-related sensory evidence alone. *p_common_*(*d*) is the weight assigned to the common-cause, fused estimate, and 1 − *p_common_*(*d*) is the weight assigned to the separate-cause, body-only estimate. For the estimate of the period, *T_fused_* = *T_body_* + *K_aud_*(*z_aud_* − *T_body_*) and *K_aud_* = 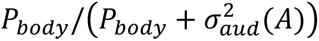, where *T_body_* and *P_body_* are the mean and variance of the step-period estimate after natural (undelayed) multisensory evidence, *z_aud_* is the amplified auditory observation, and *K_aud_* is the reliability weight assigned to it. *K_aud_* increases as auditory precision improves, so amplification enters the fused estimate only through *σ_aud_*(*A*).

This mechanism explains why non-delayed amplified feedback can reduce cadence variability: high-amplitude, zero-delay auditory feedback is both precise and likely to be self-generated, so it receives strong weight in the posterior estimate. This is consistent with the empirical finding that the no-delay, high-amplitude condition produced the clearest reduction in CV of cadence.

To account for cadence increases at larger delays, the model separated sensory estimation from action selection. After estimating the step period, the controller selected the next action under an asymmetric loss function, motivated by Bayesian decision theory (Körding, 2007; Körding & Wolpert, 2006):

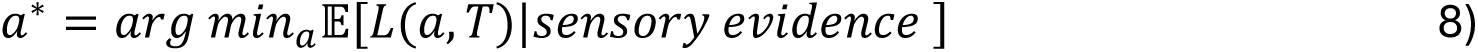

Here, *a*^∗^ is the selected action, operationalized as the next step period; *a* is a candidate action; *T* is the step-period state being acted on; *L*(*a*, *T*) is the loss associated with choosing action *a* when the relevant step-period state is *T*; and the expectation is taken with respect to the posterior belief over *T* after observing the available sensory evidence. The loss function penalized late actions more than early actions:

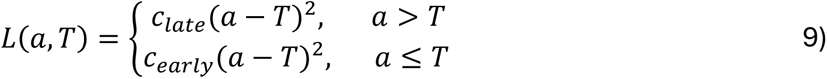

Here, *L*(*a*, *T*) is the action cost; *c_late_* is the cost coefficient for a late action; *c_early_* is the cost coefficient for an early action; *a* > *T* means that the chosen period is longer than the estimated period and is therefore treated as late; and *a* ≤ *T* means that the chosen period is shorter than or equal to the estimated period and is therefore treated as early or on time. In the model, the late-action cost increased when delayed auditory feedback was unlikely to belong to the current step:

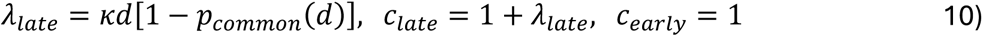

where *λ_late_* is the delay-dependent late-penalty term and *κ* is the late-penalty strength parameter. This policy term becomes larger when delay is high and common-cause attribution is low. The resulting late-penalty policy shifts the selected next step period earlier, thereby increasing cadence. This assumption is consistent with the ecological asymmetry of walking: adjusting the step too early may be suboptimal, but adjusting it too late can threaten balance, for example by failing to clear an obstacle. Thus, in the present toy model, the tendency toward early action was represented as an asymmetric action cost rather than simply as a sensory hyper-prior.

We implemented the model as a discrete-time scalar state-space process in which the step-period estimate and selected step period were updated once per simulated step. At each step, the prior estimate was first updated using undelayed body-related sensory evidence and then combined with an amplified auditory observation according to its reliability and inferred probability of arising from the current step. The resulting posterior informed action selection under the delay-dependent asymmetric late-action cost described above, and motor-execution noise scaled with within-model sensory uncertainty. The natural, unamplified baseline used a small nonzero auditory gain of 0.20. We explored the parameter space of amplification and delay in the ranges *A*=[0,1] and *d*=[0,1]. For each experimental condition, we simulated 80 trials of 600 steps and discarded the first 120 steps. Parameters were selected to illustrate the qualitative empirical patterns rather than fitted to the data. Simulation settings are in the code repository.

As Figure 7A shows, the model successfully accounted for the pattern of change in cadence observed with change in delay, namely a drop followed by an increase as delay increased. As Figure 7B shows, the model also successfully accounted for the drop in variability with increasing amplitude. Interestingly, when going beyond the range of the presently tested experimental parameters, this model predicts a monotonously increasing pattern, which is different from the periodic pattern seen with the anticipatory dynamical model, compare Figure 7 to Figures 5A and 6A. This is an exciting outcome because it means a future study can compare the two theoretical approaches on equal terms by fixing their hypotheses and exploring empirically a wide range of delays.

**Figure 7.**
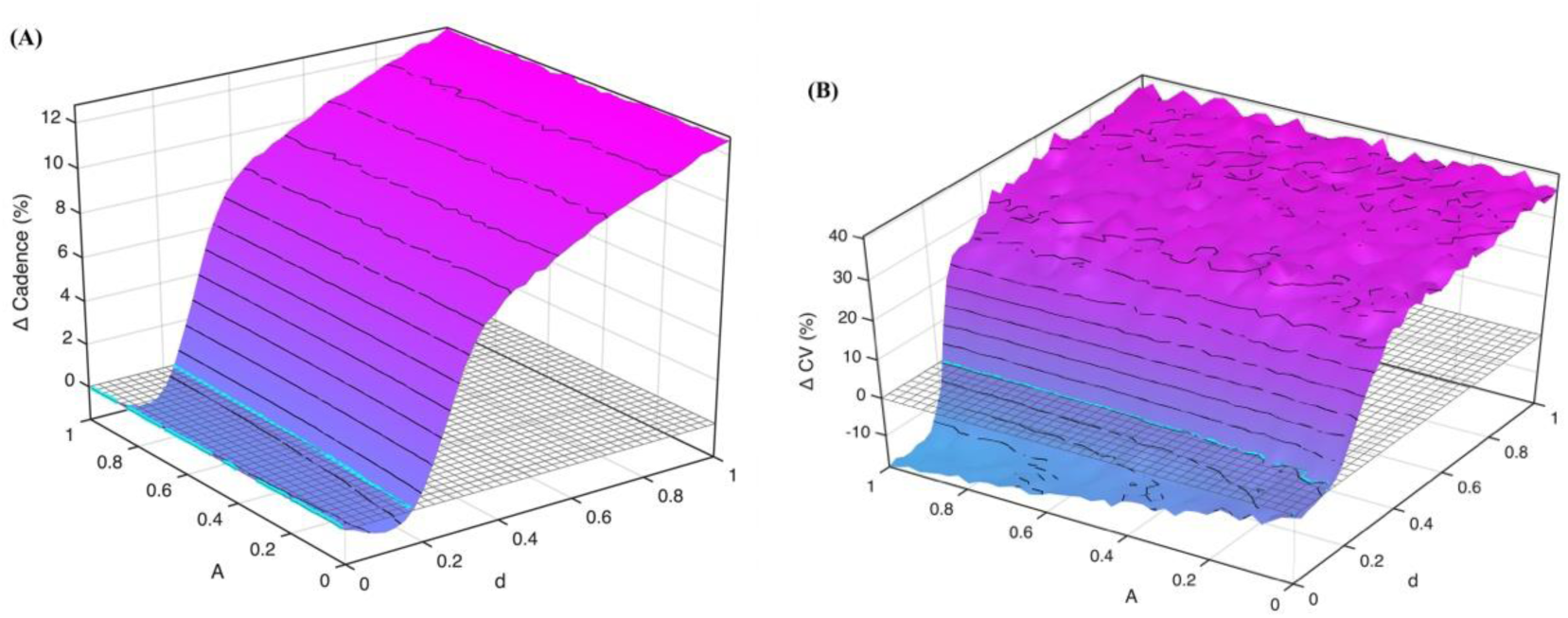
Bayesian causal-inference model of delayed and/or amplified auditory feedback. Outcomes are expressed as percent change from the natural baseline condition. A) Change in cadence. B) Change in *CV*_IOI_. Black dashed contours indicate zero change from baseline, and open circles mark the experimental delay-amplitude conditions.

### Potential of endogenous cueing

Exogenous cueing has been studied extensively to (re)train the motor rhythms involved in gait and other motor tasks in Parkinson’s disease (PD) (Cochen De Cock et al., 2018; Spaulding et al., 2013). It rests on the idea that external rhythms recruit spare pathways for target-driven gait control (Dalla Bella et al., 2017). An alternative to engaging alternative pathways, however, is to boost residual capacities for autonomous rhythm generation by amplifying its sensory component. The neural and sensory feedback from one’s own voluntary movements, also known as sensory reafference, is essential for coordinated movement and therefore it makes a major target for physical therapy (Canlı et al., 2024; X. Chen et al., 2018; Thompson et al., 2020). Although Parkinson’s disease has traditionally been viewed as a motor disorder, it also leads to sensory abnormalities and impaired sensorimotor integration. Proprioceptive deficits due to reduced sensitivity and noisy feedback contribute to miscalibration of self-generated movements and forces, and thus sensory reafference is an important therapeutic target (Lewis & Byblow, 2002; Permezel et al., 2023; Tamilselvam et al., 2025). For example, speech therapy uses feedback to train people with PD to recognize that vocalizations with increased amplitude are within normal limits and thus achieve sensory recalibration and maintenance of treatment outcomes (Fox et al., 2012). Paradoxically, DAF can also improve speech rate and intelligibility of individual patients with PD or stroked-related speech disorders such as aphasias and dysarthrias (Brendel et al., 2004; Hardy et al., 2018).

This present study is among the few to demonstrate that endogenous cueing from self-generated footstep sounds can reduce gait variability, although the generalization of AAF to populations with gait disorders remains to be evaluated. Augmenting sensory feedback to achieve endogenous cueing of gait is a novel approach grounded on the idea of self-entrainment: perceptual and motor rhythms are mutually reinforced through internal neural reciprocal connections (Cannon, 2025; Witek, 2022). Rhythm perception and sustained rhythmic production are known to engage a cortico-basal ganglia thalamo-cortical loop which involving sensory and motor areas (Balasubramaniam et al., 2021; Proksch et al., 2020). Importantly, there is a second layer of feedback that encompasses the sensory consequences of the moving body’s interaction with the environment. AAF may be reinforcing the self-entrainment of gait-related rhythms by increasing the gain and bandwidth of relevant sensory information. In simple words, AAF may enhance the coupling of sensory and motor processes from which rhythmic behavior emerges. Importantly, AAF is self-generated and as such it has the advantage of delivering ecological, variable, individualized, and continuous sounds with high action-relevance, all of which are beneficial in interventions (Dalla Bella et al., 2018; Kimijanová et al., 2024; Murgia et al., 2015, 2018; Stergiou et al., 2006; Stergiou & Decker, 2011; van der Kooij et al., 2001; Zagala et al., 2024).

There are also hearing-specific reasons for focusing on the auditory domain. Even though reduction in auditory information is a less frequently studied risk factor in safe walking, hearing loss is associated with fall risk (Lin & Ferrucci, 2012; Viljanen et al., 2009) and reduction in gait speed (L. Li et al., 2013), independently from the more obvious factors such as uneven surfaces (W. Li et al., 2006) and visual acuity (Lord, 2006). Impaired hearing can lead to poor awareness of moving objects in the spatial environment, or implicitly increase cognitive load and shared attention (Lin & Ferrucci, 2012, p. 20). Limiting auditory cues in adults with normal hearing leads to increased postural sway (Kanegaonkar et al., 2012). Importantly, hearing loss is a modifiable risk factor (Deandrea et al., 2010). Case studies report improved gait performance while wearing hearing aids (Shayman et al., 2017). In this context, it is important to verify that AAF does not interfere with gait by inducing a split task. Our study did not show evidence for increased prefrontal activity, and future work needs to confirm this in special populations such as PD where dual tasking is detrimental to gait and increases prefrontal activation (Kvist et al., 2026).

## Limitations

The main issue associated with this study was that the block number had a much stronger effect size than any of the other factors, suggesting that participants were adapting to the feedback. Condition order was randomized to address this issue, yet future studies could reduce the number of conditions and use a between-subjects design to confirm that factors did not interact with adaptation across blocks. Furthermore, the complexity of the experimental design may have limited the statistical power to detect weak effects such as possible effects of masking noise. Masking noise was intended to reduce auditory feedback, yet we could not directly determine whether it fully attenuated bone-conducted feedback. Similarly, the auditory feedback should always be seen as added on top of inherent multisensory low-delay proprioception from tactile, mechanical and vestibular stimulation. Also, because of design complexity, we could not explore the full range of delays which, as the two modeling approaches showed, may exhibit further interesting patterns of results. With respect to the hemodynamic response, we conducted an exploratory analysis that provides only a tentative idea of changes of cortical activation in a few broadly defined regions of interest. fNIRS can detect activation at relatively slow timescales and is suited for gross cortical activation patterns from sustained engagement of different sensory, motor, attentional, and cognitive modalities.

## Acknowledgements

DD received support from NIH P20GM109090 during preparation of this study. JG received support from NIH P20GM109090 and NIH 5P20GM152301-02.

## Appendix A: fNIRS mongate

**Table A1.** Source-detector channels grouped by region of interest (ROI)

| ROI | Source Label | Detector Label |
| --- | --- | --- |
| Premotor and Supplementary<br>Motor Cortex | FCC2h | FCC1h |
|  | FCC2h | FCC4h |
|  | FCC3h | FCC1h |
|  | FFC1h | FCC1h |
|  | FCC3h | FCC5h |
|  | FCC2h | FFC2h |
|  | FCC6h | FCC4h |
|  | CCP1h | FCC1h |
|  | FCC2h | CCP2h |
| Dorsolateral Prefrontal Cortex | AFF2h | AFF1h |
|  | FFC1h | AFF1h |
|  | AFF2h | FFC2h |
| Superior Temporal Gyrus | TTP8h | FTT8h |
|  | FTT7h | TTP7h |
|  | TTP8h | TPP8h |
|  | TPP7h | TTP7h |
| Primary and Association Audi-<br>tory Cortex | TTP8h | CCP6h |
|  | CCP5h | TTP7h |
| Dorsolateral Prefrontal Cortex<br>(BA46) | AFp1 | AFF5h |
|  | AFF6h | AFp2 |
|  | AFF6h | FFC6h |
|  | FFC5h | AFF5h |
|  | FFC5h | FFC3h |
|  | FFC4h | FFC6h |

**Figure A1.**
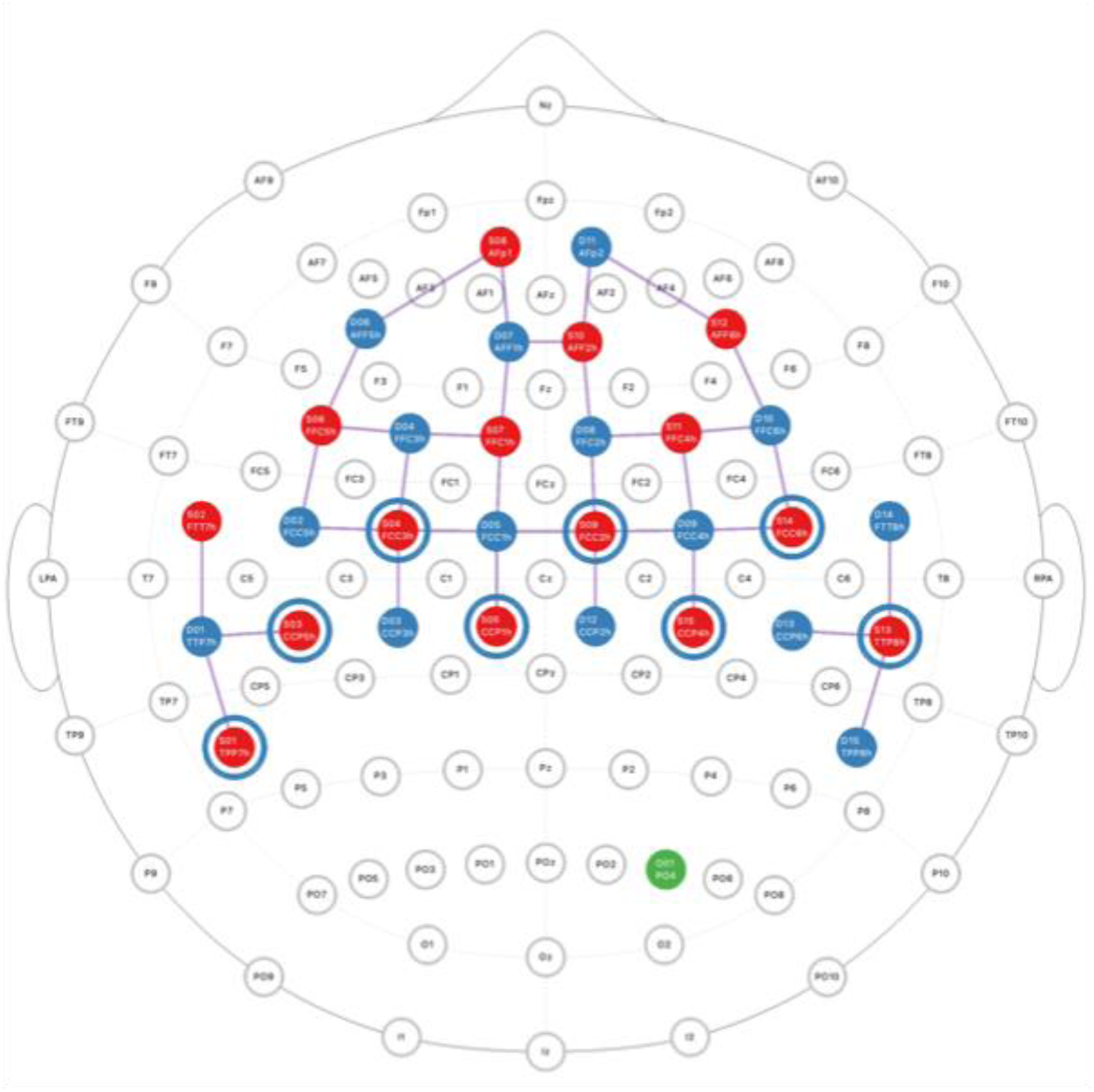
fNIRS topographical map.

## Appendix B: Descriptive statistics of gait parameters

**Table B1.** Spatiotemporal parameters of gait (mean, standard deviation, and Δ change relative to baseline) under feedback delay, amplitude, and masking manipulation.

|  | Cadence |  |  | Speed |  |  | Stride length |  |  | CV Cadence |  |  |
| --- | --- | --- | --- | --- | --- | --- | --- | --- | --- | --- | --- | --- |
| | *M* | *SD* | $\Delta$ | *M* | *SD* | $\Delta$ | *M* | *SD* | $\Delta$ | *M* | *SD* | $\Delta$ |
| Pre-test | 111.679 | 5.921 | 0.000 | 1.234 | 0.108 | 0.000 | 1.326 | 0.119 | 0.000 | 0.016 | 0.005 | 0.000 |
| Delay No, Amplitude Low | 114.676 | 6.128 | 2.747 | 1.323 | 0.169 | 7.167 | 1.382 | 0.150 | 4.187 | 0.014 | 0.005 | -9.817 |
| Delay No, Amplitude High | 114.107 | 5.899 | 2.225 | 1.311 | 0.155 | 6.195 | 1.377 | 0.144 | 3.826 | 0.013 | 0.004 | -14.813 |
| Delay Low, Amplitude Low | 113.766 | 7.022 | 1.848 | 1.309 | 0.147 | 6.039 | 1.380 | 0.132 | 4.119 | 0.016 | 0.008 | 2.832 |
| Delay Low, Amplitude High | 113.791 | 6.850 | 1.897 | 1.313 | 0.162 | 6.309 | 1.383 | 0.142 | 4.263 | 0.016 | 0.005 | -1.660 |
| Delay High, Amplitude Low | 115.442 | 6.397 | 3.436 | 1.322 | 0.164 | 6.982 | 1.372 | 0.144 | 3.365 | 0.015 | 0.005 | -7.278 |
| Delay High, Amplitude High | 115.330 | 6.378 | 3.215 | 1.319 | 0.155 | 6.975 | 1.371 | 0.135 | 3.579 | 0.015 | 0.005 | -2.641 |
| Masking Noise Low | 114.384 | 6.210 | 2.467 | 1.319 | 0.169 | 6.754 | 1.381 | 0.151 | 4.079 | 0.015 | 0.006 | -4.278 |
| Masking Noise High | 114.375 | 5.936 | 2.464 | 1.316 | 0.154 | 6.653 | 1.380 | 0.144 | 4.007 | 0.015 | 0.005 | -7.153 |
| Post-test | 115.723 | 6.367 | 3.662 | 1.353 | 0.171 | 9.514 | 1.400 | 0.151 | 5.546 | 0.015 | 0.006 | -6.514 |

## Appendix C: Statistical model tables for gait parameters

**Table C1.** Cadence.

|  | Model 0 | Model 1 | Model 2 | Model 3 | **Model 4** | Model 5 |
| --- | --- | --- | --- | --- | --- | --- |
| (Intercept) | 0.98<sup>*</sup><br>(0.43) | 0.77<br>(0.44) | 0.67<br>(0.45) | 0.82<br>(0.46) | 1.13<sup>*</sup><br>(0.46) | 1.13<sup>*</sup><br>(0.46) |
| Block |  | 0.32<sup>***</sup><br>(0.06) | 0.33<sup>***</sup><br>(0.06) | 0.32<sup>***</sup><br>(0.06) | 0.29<sup>***</sup><br>(0.06) | 0.29<sup>***</sup><br>(0.06) |
| Masking Noise |  | 0.23<sup>*</sup><br>(0.11) | 0.33<sup>**</sup><br>(0.11) | 0.36<sup>**</sup><br>(0.11) | 0.09<br>(0.11) | 0.09<br>(0.12) |
| Delay |  |  | 0.53<sup>***</sup><br>(0.09) | 0.55<sup>***</sup><br>(0.09) | -1.19<sup>***</sup><br>(0.22) | -1.19<sup>***</sup><br>(0.25) |
| Amplitude |  |  |  | -0.15<br>(0.08) | -0.09<br>(0.08) | -0.09<br>(0.09) |
| Delay<sup>2</sup> |  |  |  |  | 0.87<sup>***</sup><br>(0.10) | 0.87<sup>***</sup><br>(0.10) |
| Delay * Amplitude |  |  |  |  |  | 0.00<br>(0.09) |
| AIC | 2779.07 | 2757.96 | 2737.53 | 2735.95 | 2666.58 | 2668.57 |
| BIC | 2807.23 | 2795.51 | 2779.78 | 2782.90 | 2718.22 | 2724.91 |
| Log Likelihood | -1383.53 | -1370.98 | -1359.77 | -1357.98 | -1322.29 | -1322.29 |
| Num. obs. | 808 | 808 | 808 | 808 | 808 | 808 |
| Num. groups: Participant | 30 | 30 | 30 | 30 | 30 | 30 |
| Variance: Participant Delay | 0.35 | 0.41 | 0.16 | 0.16 | 0.16 | 0.16 |
| Variance: Participant Amplitude | 0.06 | 0.05 | 0.06 | 0.04 | 0.06 | 0.06 |
| Variance: Participant Block | 0.19 | 0.09 | 0.09 | 0.09 | 0.09 | 0.09 |
| Variance: Participant Intercept | 5.23 | 5.36 | 5.63 | 5.69 | 5.58 | 5.58 |
| Variance: Residual | 1.23 | 1.22 | 1.21 | 1.21 | 1.09 | 1.09 |

\*\*\*p < 0.001; \*\*p < 0.01; \*p < 0.05

**Table C2.**
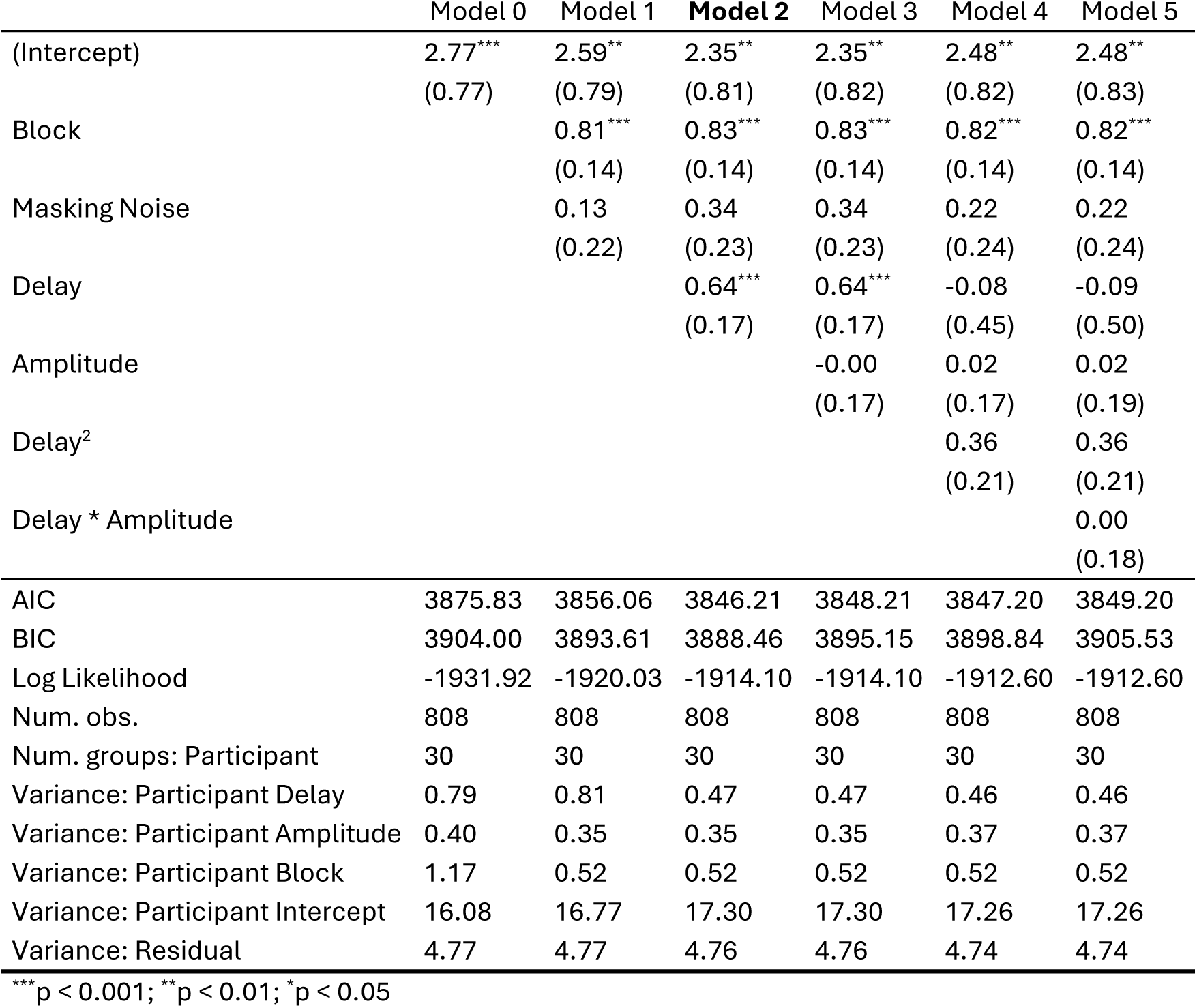
Speed.

|  | Model 0 | Model 1 | **Model 2** | Model 3 | Model 4 | Model 5 |
| --- | --- | --- | --- | --- | --- | --- |
| (Intercept) | 2.77***<br>(0.77) | 2.59**<br>(0.79) | 2.35**<br>(0.81) | 2.35**<br>(0.82) | 2.48**<br>(0.82) | 2.48**<br>(0.83) |
| Block |  | 0.81***<br>(0.14) | 0.83***<br>(0.14) | 0.83***<br>(0.14) | 0.82***<br>(0.14) | 0.82***<br>(0.14) |
| Masking Noise |  | 0.13<br>(0.22) | 0.34<br>(0.23) | 0.34<br>(0.23) | 0.22<br>(0.24) | 0.22<br>(0.24) |
| Delay |  |  | 0.64***<br>(0.17) | 0.64***<br>(0.17) | -0.08<br>(0.45) | -0.09<br>(0.50) |
| Amplitude |  |  |  | -0.00<br>(0.17) | 0.02<br>(0.17) | 0.02<br>(0.19) |
| Delay<sup>2</sup> |  |  |  |  | 0.36<br>(0.21) | 0.36<br>(0.21) |
| Delay * Amplitude |  |  |  |  |  | 0.00<br>(0.18) |
| AIC | 3875.83 | 3856.06 | 3846.21 | 3848.21 | 3847.20 | 3849.20 |
| BIC | 3904.00 | 3893.61 | 3888.46 | 3895.15 | 3898.84 | 3905.53 |
| Log Likelihood | -1931.92 | -1920.03 | -1914.10 | -1914.10 | -1912.60 | -1912.60 |
| Num. obs. | 808 | 808 | 808 | 808 | 808 | 808 |
| Num. groups: Participant | 30 | 30 | 30 | 30 | 30 | 30 |
| Variance: Participant Delay | 0.79 | 0.81 | 0.47 | 0.47 | 0.46 | 0.46 |
| Variance: Participant Amplitude | 0.40 | 0.35 | 0.35 | 0.35 | 0.37 | 0.37 |
| Variance: Participant Block | 1.17 | 0.52 | 0.52 | 0.52 | 0.52 | 0.52 |
| Variance: Participant Intercept | 16.08 | 16.77 | 17.30 | 17.30 | 17.26 | 17.26 |
| Variance: Residual | 4.77 | 4.77 | 4.76 | 4.76 | 4.74 | 4.74 |

\*\*\*p < 0.001; \*\*p < 0.01; \*p < 0.05

**Table C3.** Stride Length.

|  | Model 0 | Model 1 | Model 2 | Model 3 | **Model 4** | Model 5 |
| --- | --- | --- | --- | --- | --- | --- |
| (Intercept) | 1.77***<br>(0.38) | 1.72***<br>(0.39) | 1.68***<br>(0.40) | 1.55***<br>(0.41) | 1.35**<br>(0.41) | 1.34**<br>(0.42) |
| Block |  | 0.46***<br>(0.09) | 0.46***<br>(0.09) | 0.47***<br>(0.09) | 0.49***<br>(0.09) | 0.49***<br>(0.09) |
| Masking Noise |  | -0.05<br>(0.14) | -0.01<br>(0.14) | -0.03<br>(0.15) | 0.14<br>(0.15) | 0.14<br>(0.15) |
| Delay |  |  | 0.08<br>(0.11) | 0.06<br>(0.11) | 1.20***<br>(0.29) | 1.22***<br>(0.32) |
| Amplitude |  |  |  | 0.14<br>(0.11) | 0.11<br>(0.11) | 0.11<br>(0.12) |
| Delay<sup>2</sup> |  |  |  |  | -0.57***<br>(0.13) | -0.57***<br>(0.13) |
| Delay * Amplitude |  |  |  |  |  | -0.02<br>(0.11) |
| AIC | 3127.68 | 3111.12 | 3112.57 | 3112.85 | 3096.72 | 3098.70 |
| BIC | 3155.85 | 3148.68 | 3154.82 | 3159.79 | 3148.36 | 3155.04 |
| Log Likelihood | -1557.84 | -1547.56 | -1547.28 | -1546.42 | -1537.36 | -1537.35 |
| Num. obs. | 808 | 808 | 808 | 808 | 808 | 808 |
| Num. groups: Participant | 30 | 30 | 30 | 30 | 30 | 30 |
| Variance: participant Delay | 0.20 | 0.19 | 0.18 | 0.18 | 0.19 | 0.19 |
| Variance: participant Amplitude | 0.17 | 0.16 | 0.16 | 0.14 | 0.13 | 0.13 |
| Variance: participant Block | 0.42 | 0.21 | 0.21 | 0.21 | 0.20 | 0.20 |
| Variance: participant Intercept | 3.67 | 3.75 | 3.77 | 3.79 | 3.79 | 3.79 |
| Variance: Residual | 1.96 | 1.96 | 1.96 | 1.96 | 1.91 | 1.91 |

\*\*\*p < 0.001; \*\*p < 0.01; \*p < 0.05

**Table C4.** CV of Cadence.

|  | Model 0 | Model 1 | Model 2 | Model 3 | Model 4 | **Model 5** |
| --- | --- | --- | --- | --- | --- | --- |
| (Intercept) | -4.99**<br>(1.90) | -1.54<br>(2.73) | -3.82<br>(3.10) | 0.97<br>(3.94) | -1.79<br>(3.98) | 1.35<br>(4.23) |
| Block |  | -0.91*<br>(0.45) | -0.77<br>(0.46) | -1.02*<br>(0.48) | -0.79<br>(0.49) | -0.95<br>(0.49) |
| Masking Noise |  | -0.31<br>(2.45) | 1.36<br>(2.68) | 2.37<br>(2.72) | 4.97<br>(2.81) | 5.79*<br>(2.83) |
| Delay |  |  | 2.53<br>(1.61) | 3.16<br>(1.63) | 20.10***<br>(5.22) | 14.32*<br>(5.88) |
| Amplitude |  |  |  | -3.16*<br>(1.61) | -3.75*<br>(1.61) | -5.89**<br>(1.89) |
| Delay<sup>2</sup> |  |  |  |  | -8.54***<br>(2.50) | -8.90***<br>(2.50) |
| Delay * Amplitude |  |  |  |  |  | 4.61*<br>(2.16) |
| AIC | 7716.96 | 7713.14 | 7709.89 | 7705.26 | 7692.03 | 7686.12 |
| BIC | 7740.43 | 7746.00 | 7747.45 | 7747.51 | 7738.97 | 7737.76 |
| Log Likelihood | -3853.48 | -3849.57 | -3846.94 | -3843.63 | -3836.01 | -3832.06 |
| Num. obs. | 808 | 808 | 808 | 808 | 808 | 808 |
| Num. groups: Participant | 30 | 30 | 30 | 30 | 30 | 30 |
| Variance: Participant Delay | 25.43 | 21.65 | 19.91 | 19.60 | 21.11 | 20.96 |
| Variance: Participant Block | 1.87 | 1.60 | 1.60 | 1.64 | 1.80 | 1.88 |
| Variance: Participant Intercept | 38.79 | 44.36 | 44.11 | 44.20 | 40.48 | 37.85 |
| Variance: Residual | 753.38 | 754.17 | 753.72 | 750.68 | 739.34 | 736.08 |

\*\*\*p < 0.001; \*\*p < 0.01; \*p < 0.05

## Appendix D: Other spatiotemporal parameters of gait

**Figure D1.**
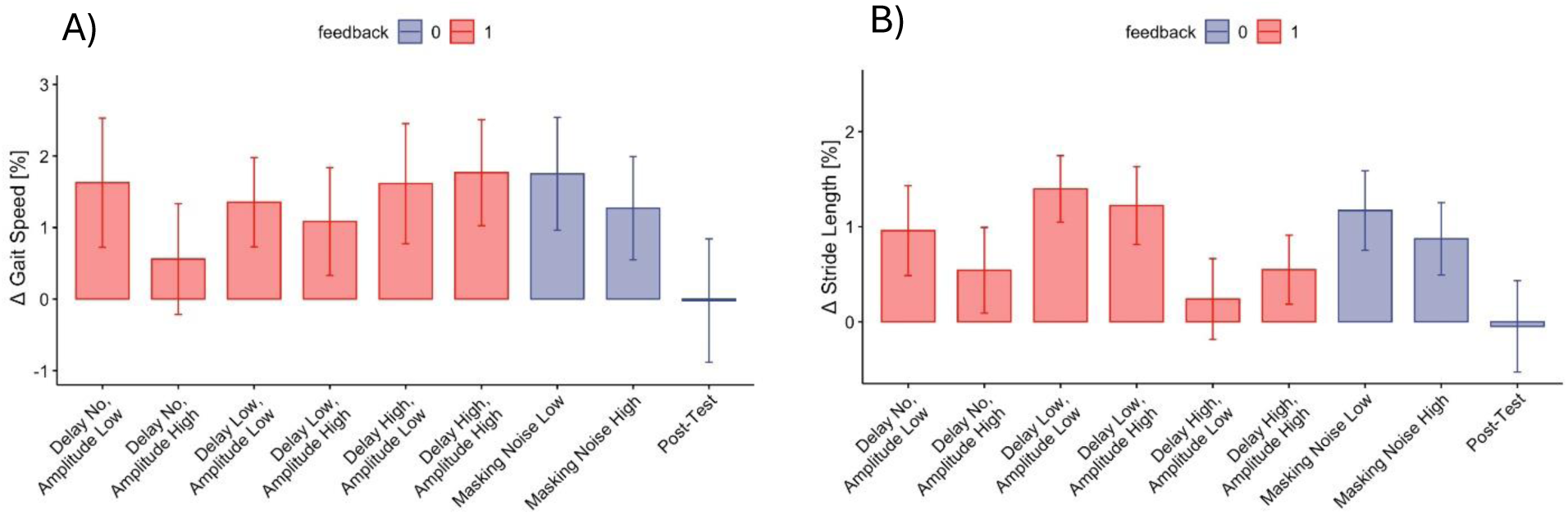
Change in spatiotemporal gait parameters in each walking condition (*M*±*SE*) relative to the baseline. The serial trend along blocks was subtracted by using the block order term in the fitted statistical model. A) Speed, averaged between left and right strides. B) Stride Length, averaged between left and right strides.

## Appendix E: Descriptive statistics of cortical hemodynamic responses

**Table E1.** Cortical activation measured in terms of the time-average of the mean hemoglobin oxygenation response function (HRF). The HRF was computed as a change relative to the pre-walking baseline period. Mean (SD) values across participants are shown per region of interest (ROI) and walking condition.

|  | dlPFC |  | PM/SMA |  | STG |  |
| --- | --- | --- | --- | --- | --- | --- |
| | $M (\times 10^{-5})$ | $SD (\times 10^{-5})$ | $M (\times 10^{-5})$ | $SD (\times 10^{-5})$ | $M (\times 10^{-5})$ | $SD (\times 10^{-5})$ |
| Pretest | 0.787 | 11.232 | -0.923 | 11.985 | 5.712 | 12.718 |
| Delay No, Amplitude Low | -0.339 | 4.659 | -1.206 | 6.884 | 8.543 | 11.275 |
| Delay No, Amplitude High | 0.341 | 11.497 | -0.027 | 11.669 | 11.873 | 15.385 |
| Delay Low, Amplitude Low | 0.193 | 5.325 | -0.984 | 5.583 | 9.239 | 9.939 |
| Delay Low, Amplitude High | -0.802 | 9.633 | -2.346 | 7.115 | 7.871 | 11.993 |
| Delay High, Amplitude Low | -2.110 | 5.230 | -3.007 | 5.814 | 6.862 | 8.888 |
| Delay High, Amplitude High | -1.117 | 5.694 | -2.077 | 6.091 | 8.752 | 10.138 |
| Masking Noise Low | -1.866 | 14.118 | 0.234 | 7.296 | 8.059 | 14.769 |
| Masking Noise High | -0.666 | 5.989 | -1.802 | 5.034 | 9.158 | 10.310 |
| Posttest | -0.450 | 6.570 | -1.433 | 6.371 | 8.632 | 11.670 |

## Footnotes

1 The pre-processed gait cycle data and subsequent modeling are available at https://github.com/dobri/dafw_kinematics_share.

2 Available at https://github.com/dobri/dafw_kinematics_share.

3 Available at https://github.com/dobri/dafw_kinematics_share.

## References

Aschersleben, G. (2002). Temporal control of movements in sensorimotor synchronization. Brain and Cognition, 48(1), 66–79. 10.1006/brcg.2001.1304

Aschersleben, G., & Prinz, W. (1997). Delayed Auditory Feedback in Synchronization. Journal of Motor Behavior, 29(1), 35–46. 10.1080/00222899709603468

Baker, K., Rochester, L., & Nieuwboer, A. (2007). The Immediate Effect of Attentional, Auditory, and a Combined Cue Strategy on Gait During Single and Dual Tasks in Parkinson’s Disease. Archives of Physical Medicine and Rehabilitation, 88(12), 1593–1600. 10.1016/j.apmr.2007.07.026

Balasubramaniam, R., Haegens, S., Jazayeri, M., Merchant, H., Sternad, D., & Song, J.-H. (2021). Neural Encoding and Representation of Time for Sensorimotor Control and Learning. Journal of Neuroscience, 41(5), 866–872. 10.1523/JNEUROSCI.1652-20.2020

Bates, D. M. (2010). lme4: Mixed-effects modeling with R. Springer.

Brendel, B., Lowit, A., & Howell, P. (2004). The effects of delayed and frequency shifted feedback on speakers with Parkinson’s Disease. Journal of Medical Speech-Language Pathology, 12, 131–138.

Canlı, K., Van Oijen, J., Van Oosterwijck, J., Meeus, M., Van Oosterwijck, S., & De Meulemeester, K. (2024). Influence of sensory retraining on cortical reorganization in peripheral neuropathy: A systematic review. PM & R: The Journal of Injury, Function, and Rehabilitation, 16(8), 888–907. 10.1002/pmrj.13126

Cannon, J. (2021). Expectancy-based rhythmic entrainment as continuous Bayesian inference. PLOS Computational Biology, 17(6), e1009025. 10.1371/journal.pcbi.1009025

Cannon, J. (2025). Marching to Your Own Beat: Self-entrainment as a Missing Link Between Rhythm Perception and Synchronization. Music Perception, 43(1), 49–60. 10.1525/mp.2025.2427447

Chen, J. L., Penhune, V. B., & Zatorre, R. J. (2009). The Role of Auditory and Premotor Cortex in Sensorimotor Transformations. Annals of the New York Academy of Sciences, 1169(1), 15–34. 10.1111/j.1749-6632.2009.04556.x

Chen, X., Liu, F., Yan, Z., Cheng, S., Liu, X., Li, H., & Li, Z. (2018). Therapeutic effects of sensory input training on motor function rehabilitation after stroke. Medicine, 97(48), e13387. 10.1097/MD.0000000000013387

Cochen De Cock, V., Dotov, D. G., Ihalainen, P., Bégel, V., Galtier, F., Lebrun, C., Picot, M. C., Driss, V., Landragin, N., Geny, C., Bardy, B., & Dalla Bella, S. (2018). Rhythmic abilities and musical training in Parkinson’s disease: Do they help? Npj Parkinson’s Disease, 4(1), Article 1. 10.1038/s41531-018-0043-7

Cornwell, T., Woodward, J., Wu, M., Jackson, B., Souza, P., Siegel, J., Dhar, S., & Gordon, K. E. (2020). Walking With Ears: Altered Auditory Feedback Impacts Gait Step Length in Older Adults. Frontiers in Sports and Active Living, 2. https://www.frontiersin.org/articles/10.3389/fspor.2020.00038

D’Adamo, A., Bevilacqua, F., Matamala-Gómez, M., & Tajadura-Jiménez, A. (2026). SoniFootsteps: Movement-Triggered Footstep Sounds to Modulate Body-Weight Perception, Gait and Emotion. Proceedings of the 10th International Conference on Movement and Computing, MOCO’26, 1–12. 10.1145/3802842.3802882

Dalla Bella, S., Benoit, C.-E. E., Farrugia, N., Keller, P. E., Obrig, H., Mainka, S., & Kotz, S. A. (2017). Gait improvement via rhythmic stimulation in Parkinson’s disease is linked to rhythmic skills. Scientific Reports, 7(October 2015), 1–11. 10.1038/srep42005

Dalla Bella, S., Dotov, D., Bardy, B., & de Cock, V. C. (2018). Individualization of music-based rhythmic auditory cueing in Parkinson’s disease. Annals of the New York Academy of Sciences, 1423(1), 308–317. 10.1111/nyas.13859

Deandrea, S., Lucenteforte, E., Bravi, F., Foschi, R., La Vecchia, C., & Negri, E. (2010). Risk factors for falls in community-dwelling older people: A systematic review and meta-analysis. Epidemiology (Cambridge, Mass.), 21(5), 658–668. 10.1097/EDE.0b013e3181e89905

Demos, A. P., Layeghi, H., Wanderley, M. M., & Palmer, C. (2019). Staying Together: A Bidirectional Delay–Coupled Approach to Joint Action. Cognitive Science, 43(8), e12766. 10.1111/cogs.12766

Dubois, D. M. (2003). Mathematical Foundations of Discrete and Functional Systems with Strong and Weak Anticipations. In M. V. Butz, O. Sigaud, & P. Gérard (Eds.), Anticipatory Behavior in Adaptive Learning Systems: Foundations, Theories, and Systems (pp. 110–132). Springer. 10.1007/978-3-540-45002-3_7

Ermentrout, B. (1991). An adaptive model for synchrony in the firefly Pteroptyx malaccae. Journal of Mathematical Biology, 29(6), 571–585. 10.1007/BF00164052

Ernst, M. O., & Banks, M. S. (2002). Humans integrate visual and haptic information in a statistically optimal fashion. Nature, 415(6870), 429–433. 10.1038/415429a

Fox, C., Ebersbach, G., Ramig, L., & Sapir, S. (2012). LSVT LOUD and LSVT BIG: Behavioral Treatment Programs for Speech and Body Movement in Parkinson Disease. Parkinson’s Disease, 2012(1), 391946. 10.1155/2012/391946

Grahn, J. A., & Rowe, J. B. (2009). Feeling the Beat: Premotor and Striatal Interactions in Musicians and Nonmusicians during Beat Perception. Journal of Neuroscience, 29(23), 7540–7548. 10.1523/JNEUROSCI.2018-08.2009

Grover, F. M., Riehm, C., Silva, P. L., Lorenz, T., & Riley, M. A. (2021). Grip force anticipation of nonlinear, underactuated load force. Journal of Neurophysiology, 125(5), 1647–1662. 10.1152/jn.00616.2020

Harding, E. E., Kim, J. C., Demos, A. P., Roman, I. R., Tichko, P., Palmer, C., & Large, E. W. (2025). Musical neurodynamics. Nature Reviews Neuroscience, 26(5), 293–307. 10.1038/s41583-025-00915-4

Hardy, C. J. D., Bond, R. L., Jaisin, K., Marshall, C. R., Russell, L. L., Dick, K., Crutch, S. J., Rohrer, J. D., & Warren, J. D. (2018). Sensitivity of Speech Output to Delayed Auditory Feedback in Primary Progressive Aphasias. Frontiers in Neurology, 9. 10.3389/fneur.2018.00894

Hashimoto, Y., & Sakai, K. L. (2003). Brain activations during conscious self-monitoring of speech production with delayed auditory feedback: An fMRI study. Human Brain Mapping, 20(1), 22–28. 10.1002/hbm.10119

Heggli, O. A., Cabral, J., Konvalinka, I., Vuust, P., & Kringelbach, M. L. (2019). A Kuramoto model of self-other integration across interpersonal synchronization strategies. PLOS Computational Biology, 15(10), e1007422. 10.1371/journal.pcbi.1007422

Hermann, T., Hunt, A., & Neuhoff, J. G. (2011). The Sonification Handbook. In The Sonification Handbook. Logos Verlag. 10.1017/CBO9781107415324.004

Hickok, G., & Poeppel, D. (2007). The cortical organization of speech processing. Nature Reviews Neuroscience, 8(5), 393–402. 10.1038/nrn2113

Huppert, T. J., Diamond, S. G., Franceschini, M. A., & Boas, D. A. (2009). HomER: A review of time-series analysis methods for near-infrared spectroscopy of the brain. Applied Optics, 48(10), D280–D298. 10.1364/AO.48.00D280

Kalinowski, J., & Stuart, A. (1996). Stuttering amelioration at various auditory feedback delays and speech rates. *European Journal of Disorders of Communication: The Journal of the College of Speech and Language Therapists*, London, 31(3), 259–269. 10.3109/13682829609033157

Kanegaonkar, R. G., Amin, K., & Clarke, M. (2012). The contribution of hearing to normal balance. The Journal of Laryngology & Otology, 126(10), 984–988. 10.1017/S002221511200179X

Kimijanová, J., Svoboda, Z., & Han, J. (2024). Editorial: Sensory control of posture and gait: integration and mechanisms to maintain balance during different sensory conditions. Frontiers in Human Neuroscience, 18. 10.3389/fnhum.2024.1378599

Kinder, K. T., Heim, H. L. R., Parker, J., Lowery, K., McCraw, A., Eddings, R. N., Defenderfer, J., Sullivan, J., & Buss, A. T. (2022). Systematic review of fNIRS studies reveals inconsistent chromophore data reporting practices. Neurophotonics, 9(4), 040601. 10.1117/1.NPh.9.4.040601

Koenraadt, K. L. M., Roelofsen, E. G. J., Duysens, J., & Keijsers, N. L. W. (2014). Cortical control of normal gait and precision stepping: An fNIRS study. NeuroImage, 85 Pt 1, 415–422. 10.1016/j.neuroimage.2013.04.070

Körding, K. (2007). Decision Theory: What “Should” the Nervous System Do? Science, 318(5850), 606–610. 10.1126/science.1142998

Körding, K., Beierholm, U., Ma, W. J., Quartz, S., Tenenbaum, J. B., & Shams, L. (2007). Causal Inference in Multisensory Perception. PLOS ONE, 2(9), e943. 10.1371/journal.pone.0000943

Körding, K., & Wolpert, D. M. (2004). Bayesian integration in sensorimotor learning. Nature, 427(6971), 244–247. 10.1038/nature02169

Körding, K., & Wolpert, D. M. (2006). Bayesian decision theory in sensorimotor control. Trends in Cognitive Sciences, 10(7), 319–326. 10.1016/j.tics.2006.05.003

Krelove, L. M., Machula, A., Sergio, L. E., & Mochizuki, G. (2025). Prefrontal cortex activity during virtual obstacle avoidance and distracted walking: A methodological proof of concept using augmented reality and functional near-infrared spectroscopy. PLOS One, 20(10), e0333622. 10.1371/journal.pone.0333622

Kvist, A., Bezuidenhout, L., Johansson, H., Albrecht, F., Ekman, U., Conradsson, D. M., & Franzén, E. (2023). Using functional near-infrared spectroscopy to measure prefrontal cortex activity during dual-task walking and navigated walking: A feasibility study. Brain and Behavior, 13(4), e2948. 10.1002/brb3.2948

Kvist, A., Peterson, D. S., Bezuidenhout, L., Johansson, H., Albrecht, F., Ekman, U., Conradsson, D. M., & Franzén, E. (2026). Exploring gait automaticity and prefrontal brain activity during single and dual-task walking in aging and Parkinson’s disease. Journal of NeuroEngineering and Rehabilitation, 23(1), 41. 10.1186/s12984-025-01864-w

Large, E. W., Roman, I., Kim, J. C., Cannon, J., Pazdera, J. K., Trainor, L. J., Rinzel, J., & Bose, A. (2023). Dynamic models for musical rhythm perception and coordination. Frontiers in Computational Neuroscience, 17. 10.3389/fncom.2023.1151895

Lewis, G. N., & Byblow, W. D. (2002). Altered sensorimotor integration in Parkinson’s disease. Brain: A Journal of Neurology, 125(Pt 9), 2089–2099. 10.1093/brain/awf200

Li, L., Simonsick, E. M., Ferrucci, L., & Lin, F. R. (2013). Hearing loss and gait speed among older adults in the United States. Gait & Posture, 38(1), 25–29. 10.1016/j.gaitpost.2012.10.006

Li, W., Keegan, T. H. M., Sternfeld, B., Sidney, S., Quesenberry, C. P., & Kelsey, J. L. (2006). Outdoor Falls Among Middle-Aged and Older Adults: A Neglected Public Health Problem. American Journal of Public Health, 96(7), 1192–1200. 10.2105/AJPH.2005.083055

Lin, F. R., & Ferrucci, L. (2012). Hearing Loss and Falls Among Older Adults in the United States. Archives of Internal Medicine, 172(4), 369–371. 10.1001/archinternmed.2011.728

Lord, S. R. (2006). Visual risk factors for falls in older people. Age and Ageing, 35(suppl_2), ii42–ii45. 10.1093/ageing/afl085

Menzer, F., Brooks, A., Halje, P., Faller, C., Vetterli, M., & Blanke, O. (2010). Feeling in control of your footsteps: Conscious gait monitoring and the auditory consequences of footsteps. Cognitive Neuroscience, 1, 184–192. 10.1080/17588921003743581

Merchant, H., Grahn, J., Trainor, L., Rohrmeier, M., & Fitch, W. T. (2015). Finding the beat: A neural perspective across humans and non-human primates. Philosophical Transactions of the Royal Society B: Biological Sciences, 370(1664), 20140093. 10.1098/rstb.2014.0093

Mirasso, C. R., Carelli, P. V., Pereira, T., Matias, F. S., & Copelli, M. (2017). Anticipated and zero-lag synchronization in motifs of delay-coupled systems. Chaos: An Interdisciplinary Journal of Nonlinear Science, 27(11), 114305. 10.1063/1.5006932

Murgia, M., Pili, R., Corona, F., Sors, F., Agostini, T. A., Bernardis, P., Casula, C., Cossu, G., Guicciardi, M., & Pau, M. (2018). The Use of Footstep Sounds as Rhythmic Auditory Stimulation for Gait Rehabilitation in Parkinson’s Disease: A Randomized Controlled Trial. Frontiers in Neurology, 9. 10.3389/fneur.2018.00348

Murgia, M., Santoro, I., Tamburini, G., Prpic, V., Sors, F., Galmonte, A., & Agostini, T. (2015). Ecological sounds affect breath duration more than artificial sounds. Psychological Research. 10.1007/s00426-015-0647-z

Nourski, K. V., Reale, R. A., Oya, H., Kawasaki, H., Kovach, C. K., Chen, H., Howard, M. A., & Brugge, J. F. (2009). Temporal Envelope of Time-Compressed Speech Represented in the Human Auditory Cortex. Journal of Neuroscience, 29(49), 15564–15574. 10.1523/JNEUROSCI.3065-09.2009

Obleser, J., Zimmermann, J., Van Meter, J., & Rauschecker, J. P. (2007). Multiple Stages of Auditory Speech Perception Reflected in Event-Related fMRI. Cerebral Cortex, 17(10), 2251–2257. 10.1093/cercor/bhl133

Pang, T. Y., Cheng, C.-T., Feltham, F., Rahman, A., McCarney, L., & Rodriguez, C. Q. (2025). Feasibility of IMU-Based Wearable Sonification: Toward Personalized, Real-Time Gait Monitoring and Rehabilitation. Biosensors, 15(10), 698. 10.3390/bios15100698

Pang, T. Y., Feltham, F., & Cheng, C.-T. (2025). Engineering Auditory Cues for Gait Modulation: Effects of Continuous and Discrete Sound Features. Eng, 6(12), 349. 10.3390/eng6120349

Park, S. H., & Lee, P. J. (2017). Effects of floor impact noise on psychophysiological responses. Building and Environment, 116, 173–181. 10.1016/j.buildenv.2017.02.005

Permezel, F., Alty, J., Harding, I. H., & Thyagarajan, D. (2023). Brain Networks Involved in Sensory Perception in Parkinson’s Disease: A Scoping Review. Brain Sciences, 13(11), 1552. 10.3390/brainsci13111552

Perrey, S. (2014). Possibilities for examining the neural control of gait in humans with fNIRS. Frontiers in Physiology, 5, 204. 10.3389/fphys.2014.00204

Pfordresher, P. Q., & Dalla Bella, S. (2011). Delayed auditory feedback and movement. Journal of Experimental Psychology: Human Perception and Performance, 37(2), 566–579. 10.1037/a0021487

Pirini, M., Mancini, M., Farella, E., & Chiari, L. (2011). EEG correlates of postural audio-biofeedback. *Human Movement Science*, Basic Mechanisms Underlying Balance Control under Static and Dynamic Conditions, 30(2), 249–261. 10.1016/j.humov.2010.05.016

Ploughman, M., Shears, J., Quinton, S., Flight, C., O’brien, M., MacCallum, P., Kirkland, M. C., & Byrne, J. M. (2018). Therapists’ cues influence lower limb muscle activation and kinematics during gait training in subacute stroke. Disability and Rehabilitation, 40(26), 3156–3163. 10.1080/09638288.2017.1380720

Proksch, S., Comstock, D. C., Médé, B., Pabst, A., & Balasubramaniam, R. (2020). Motor and Predictive Processes in Auditory Beat and Rhythm Perception. Frontiers in Human Neuroscience, 14. https://www.frontiersin.org/articles/10.3389/fnhum.2020.578546

Rahman, T. T., Polskaia, N., St-Amant, G., Salzman, T., Vallejo, D. T., Lajoie, Y., & Fraser, S. A. (2021). An fNIRS Investigation of Discrete and Continuous Cognitive Demands During Dual-Task Walking in Young Adults. Frontiers in Human Neuroscience, 15, 711054. 10.3389/fnhum.2021.711054

Raudenbush, S. W., & Bryk, A. S. (2002). Hierarchical Linear Models: Applications and Data Analysis Methods. SAGE.

Rauschecker, J. P., & Scott, S. K. (2009). Maps and streams in the auditory cortex: Nonhuman primates illuminate human speech processing. Nature Neuroscience, 12(6), 718–724. 10.1038/nn.2331

Ready, E. A., Holmes, J. D., & Grahn, J. A. (2022). Gait in younger and older adults during rhythmic auditory stimulation is influenced by groove, familiarity, beat perception, and synchronization demands. Human Movement Science, 84, 102972. 10.1016/j.humov.2022.102972

Reznik, D., Henkin, Y., Schadel, N., & Mukamel, R. (2014). Lateralized enhancement of auditory cortex activity and increased sensitivity to self-generated sounds. Nature Communications, 5(1), 4059. 10.1038/ncomms5059

Rochester, L., Hetherington, V., Jones, D., Nieuwboer, A., Willems, A.-M., Kwakkel, G., & Van Wegen, E. (2005). The Effect of External Rhythmic Cues (Auditory and Visual) on Walking During a Functional Task in Homes of People With Parkinson’s Disease. Archives of Physical Medicine and Rehabilitation, 86(5), 999–1006. 10.1016/j.apmr.2004.10.040

Rohe, T., & Noppeney, U. (2015). Cortical Hierarchies Perform Bayesian Causal Inference in Multisensory Perception. PLOS Biology, 13(2), e1002073. 10.1371/journal.pbio.1002073

Roman, I. R., Washburn, A., Large, E. W., Chafe, C., & Fujioka, T. (2019). Delayed feedback embedded in perception-action coordination cycles results in anticipation behavior during synchronized rhythmic action: A dynamical systems approach. PLoS Computational Biology, 15(10). 10.1371/journal.pcbi.1007371

Ross, B., Barat, M., & Fujioka, T. (2017). Sound-Making Actions Lead to Immediate Plastic Changes of Neuromagnetic Evoked Responses and Induced β-Band Oscillations during Perception. Journal of Neuroscience, 37(24), 5948–5959. 10.1523/JNEUROSCI.3613-16.2017

Roytman, S., Paalanen, R., Carli, G., Marusic, U., Kanel, P., van Laar, T., & Bohnen, N. I. (2025). Multisensory mechanisms of gait and balance in Parkinson’s disease: An integrative review. Neural Regeneration Research, 20(1), 82. 10.4103/NRR.NRR-D-23-01484

Schmitz, G., Kroeger, D., Effenberg, A. O., & Moritzwinkel, A. (2014). A mobile sonification system for stroke rehabilitation Institute of Sports Science, Leibniz University Hannover,.

Schneider, D. M., & Mooney, R. (2018). How Movement Modulates Hearing. Annual Review of Neuroscience, 41(1), 553–572. 10.1146/annurev-neuro-072116-031215

Schneider, D. M., Sundararajan, J., & Mooney, R. (2018). A cortical filter that learns to suppress the acoustic consequences of movement. Nature, 561(7723), Article 7723. 10.1038/s41586-018-0520-5

Shayman, C. S., Earhart, G. M., & Hullar, T. E. (2017). Improvements in gait with hearing aids and cochlear implants. *Otology & Neurotology : Official Publication of the American Otological Society*, American Neurotology Society [and] European Academy of Otology and Neurotology, 38(4), 484–486. 10.1097/MAO.0000000000001360

Shin, J., & Chung, Y. (2022). The effects of treadmill training with visual feedback and rhythmic auditory cue on gait and balance in chronic stroke patients: A randomized controlled trial. NeuroRehabilitation, 51(3), 443–453. 10.3233/NRE-220099

Singer, J. D., & Willett, J. B. (2003). Applied Longitudinal Data Analysis: Modeling Change and Event Occurrence. Oxford University Press.

Spaulding, S. J., Barber, B., Colby, M., Cormack, B., Mick, T., & Jenkins, M. E. (2013). Cueing and Gait Improvement Among People With Parkinson’s Disease: A Meta-Analysis. Archives of Physical Medicine and Rehabilitation, 94(3), 562–570. 10.1016/j.apmr.2012.10.026

Spencer, J., Wolf, S. L., & Kesar, T. M. (2021). Biofeedback for Post-stroke Gait Retraining: A Review of Current Evidence and Future Research Directions in the Context of Emerging Technologies. Frontiers in Neurology, 12. 10.3389/fneur.2021.637199

Steen, M. C. (Marieke) van der, & Keller, P. E. (2013). The ADaptation and Anticipation Model (ADAM) of sensorimotor synchronization. Frontiers in Human Neuroscience, 7, 253. 10.3389/fnhum.2013.00253

Stepp, N. (2009). Anticipation in feedback-delayed manual tracking of a chaotic oscillator. Experimental Brain Research, 198(4), 521–525. 10.1007/s00221-009-1940-0

Stepp, N., & Turvey, M. T. (2015). The Muddle of Anticipation. Ecological Psychology, 27(2), 103–126. 10.1080/10407413.2015.1027123

Stergiou, N., & Decker, L. M. (2011). Human movement variability, nonlinear dynamics, and pathology: Is there a connection? Human Movement Science, EWOMS 200S: The European Workshop on Movement Science, 30(5), 869–888. 10.1016/j.humov.2011.06.002

Stergiou, N., Harbourne, R. T., & Cavanaugh, J. T. (2006). Optimal Movement Variability: A New Theoretical Perspective for Neurologic Physical Therapy. Journal of Neurologic Physical Therapy, 30(3), 120–129.

Stuart, A., Kalinowski, J., Rastatter, M. P., & Lynch, K. (2002). Effect of delayed auditory feedback on normal speakers at two speech rates. The Journal of the Acoustical Society of America, 111(5), 2237–2241. 10.1121/1.1466868

Stuart, A., Kalinowski, J., Rastatter, M. P., Saltuklaroglu, T., & Dayalu, V. (2004). Investigations of the impact of altered auditory feedback in-the-ear devices on the speech of people who stutter: Initial fitting and 4-month follow-up. International Journal of Language & Communication Disorders, 39(1), 93–113. 10.1080/13682820310001616976

Tachibana, A., Noah, J. A., Bronner, S., Ono, Y., & Onozuka, M. (2011). Parietal and temporal activity during a multimodal dance video game: An fNIRS study. Neuroscience Letters, 503(2), 125–130. 10.1016/j.neulet.2011.08.023

Tajadura-Jiménez, A., Basia, M., Deroy, O., Fairhurst, M., Marquardt, N., & Bianchi-Berthouze, N. (2015). As Light as your Footsteps. Proceedings of the 33rd Annual ACM Conference on Human Factors in Computing Systems - CHI’ 15, 2943–2952. 10.1145/2702123.2702374

Tamilselvam, Y. K., Ganguly, J., Jog, M. S., & Patel, R. V. (2025). Sensorimotor Integration: A Review of Neural and Computational Models and the Impact of Parkinson’s Disease. IEEE Transactions on Cognitive and Developmental Systems, 17(1), 3–21. 10.1109/TCDS.2024.3520976

Taylor, D., Ott, E., & Restrepo, J. G. (2010). Spontaneous synchronization of coupled oscillator systems with frequency adaptation. Physical Review E, 81(4), 046214. 10.1103/PhysRevE.81.046214

Thaut, M. H., McIntosh, G. C., & Rice, R. R. (1997). Rhythmic facilitation of gait training in hemiparetic stroke rehabilitation. Journal of the Neurological Sciences, 151(2), 207–212. 10.1016/S0022-510X(97)00146-9

Thaut, M. H., McIntosh, G. C., Rice, R. R., Miller, R. A., Rathbun, J., & Brault, J. M. (1996). Rhythmic auditory stimulation in gait training for Parkinson’s disease patients. Movement Disorders, 11(2), 193–200. 10.1002/mds.870110213

Thompson, L. A., Savadkoohi, M., de Paiva, G. V., Augusto Renno Brusamolin, J., Guise, J., Suh, P., & Guerrero, P. S. (2020). Sensory integration training improves balance in older individuals. Annual International Conference of the IEEE Engineering in Medicine and Biology Society. IEEE Engineering in Medicine and Biology Society. Annual International Conference, 2020, 3811–3814. 10.1109/EMBC44109.2020.9175715

van der Kooij, H., Jacobs, R., Koopman, B., & van der Helm, F. (2001). An adaptive model of sensory integration in a dynamic environment applied to human stance control. Biological Cybernetics, 84(2), 103–115. 10.1007/s004220000196

van Elk, M., Salomon, R., Kannape, O., & Blanke, O. (2014). Suppression of the N1 auditory evoked potential for sounds generated by the upper and lower limbs. Biological Psychology, 102, 108–117. 10.1016/j.biopsycho.2014.06.007

Viljanen, A., Kaprio, J., Pyykkö, I., Sorri, M., Pajala, S., Kauppinen, M., Koskenvuo, M., & Rantanen, T. (2009). Hearing as a Predictor of Falls and Postural Balance in Older Female Twins. The Journals of Gerontology: Series A, 64A(2), 312–317. 10.1093/gerona/gln015

Vitorio, R., Stuart, S., Gobbi, L. T. B., Rochester, L., Alcock, L., & Pantall, A. (2018). Reduced Gait Variability and Enhanced Brain Activity in Older Adults With Auditory Cues: A Functional Near-Infrared Spectroscopy Study. Neurorehabilitation and Neural Repair, 32(11), 976–987. 10.1177/1545968318805159

Voss, H. U. (2000). Anticipating chaotic synchronization. Physical Review E, 61(5), 5115–5119. 10.1103/PhysRevE.61.5115

Voss, H. U., & Stepp, N. (2016). A negative group delay model for feedback-delayed manual tracking performance. Journal of Computational Neuroscience, 41(3), 295–304. 10.1007/s10827-016-0618-4

Vuust, P., & Witek, M. A. G. (2014). Rhythmic complexity and predictive coding: A novel approach to modeling rhythm and meter perception in music. Frontiers in Psychology, 5. https://www.frontiersin.org/articles/10.3389/fpsyg.2014.01111

Wall, C., McMeekin, P., Walker, R., Hetherington, V., Graham, L., & Godfrey, A. (2024). Sonification for Personalised Gait Intervention. Sensors, 24(1), Article 1. 10.3390/s24010065

Washburn, A., Kallen, R. W., Coey, C. A., Shockley, K., & Richardson, M. J. (2015). Harmony from Chaos? Perceptual-Motor Delays Enhance Behavioral Anticipation in Social Interaction. Journal of Experimental Psychology: Human Perception and Performance, 41(4), 1166–1177. 10.1037/xhp0000080

Wilson, T. W., Kurz, M. J., & Arpin, D. J. (2014). Functional specialization within the supplementary motor area: A fNIRS study of bimanual coordination. NeuroImage, Celebrating 20 Years of Functional Near Infrared Spectroscopy (fNIRS), 85, 445–450. 10.1016/j.neuroimage.2013.04.112

Witek, M. (2022). The mind is a DJ: Rhythmic entrainment in beatmatching and embodied temporal processing. In M. Doffman, E. Payne, & T. Young (Eds.), The Oxford Handbook of Time in Music (pp. 234–252). Oxford University Press. 10.1093/oxfordhb/9780190947279.013.13

Wolpert, D. M., Ghahramani, Z., & Jordan, M. I. (1995). An Internal Model for Sensorimotor Integration. Science, 269(5232), 1880–1882. 10.1126/science.7569931

Yates, A. J. (1963). Delayed auditory feedback. Psychological Bulletin, 60(3), 213–232. 10.1037/h0044155

Yücel, M. A., Lühmann, A. v, Scholkmann, F., Gervain, J., Dan, I., Ayaz, H., Boas, D., Cooper, R. J., Culver, J., Elwell, C. E., Eggebrecht, A., Franceschini, M. A., Grova, C., Homae, F., Lesage, F., Obrig, H., Tachtsidis, I., Tak, S., Tong, Y., … Wolf, M. (2021). Best practices for fNIRS publications. Neurophotonics, 8(1), 012101. 10.1117/1.NPh.8.1.012101

Zagala, A., Foster, N. E. V., Vugt, F. T. van, Maso, F. D., & Dalla Bella, S. (2024). *The Ramp protocol: Uncovering individual differences in walking to an auditory beat using TeensyStep* (p. 2024.03.07.583713). bioRxiv. 10.1101/2024.03.07.583713

Zatorre, R. J., Chen, J. L., & Penhune, V. B. (2007). When the brain plays music: Auditory– motor interactions in music perception and production. Nature Reviews Neuroscience, 8(7), 547–558. 10.1038/nrn2152

Zimeo Morais, G. A., Balardin, J. B., & Sato, J. R. (2018). fNIRS Optodes’ Location Decider (fOLD): A toolbox for probe arrangement guided by brain regions-of-interest. Scientific Reports, 8(1), 3341. 10.1038/s41598-018-21716-z

